# Genomes of Thaumarchaeota from deep sea sediments reveal specific adaptations of three independently evolved lineages

**DOI:** 10.1101/2020.06.24.168906

**Authors:** Melina Kerou, Rafael I. Ponce-Toledo, Rui Zhao, Sophie S. Abby, Miho Hirai, Hidetaka Nomaki, Yoshihiro Takaki, Takuro Nunoura, Steffen L. Jørgensen, Christa Schleper

## Abstract

Marine sediments represent a vast habitat for complex microbiomes. Among these, ammonia oxidizing archaea (AOA) of the phylum Thaumarchaeota are one of the most common, yet little explored inhabitants, that seem extraordinarily well adapted to the harsh conditions of the subsurface biosphere. We present 11 metagenome-assembled genomes of the most abundant AOA clades from sediment cores obtained from the Atlantic Mid-Ocean ridge flanks and Pacific abyssal plains. Their phylogenomic placement reveals three independently evolved clades within the order *Ca.* Nitrosopumilales, of which no cultured representative is known yet. In addition to the gene sets for ammonia oxidation and carbon fixation known from other AOA, all genomes encode an extended capacity for the conversion of fermentation products that can be channeled into the central carbon metabolism, as well as uptake of amino acids probably for protein maintenance or as an ammonia source. Two lineages encode an additional (V-type) ATPase and a large repertoire of gene repair systems that may allow to overcome challenges of high hydrostatic pressure. We suggest that the adaptive radiation of AOA into marine sediments occurred more than once in evolution and resulted in three distinct lineages with particular adaptations to this extremely energy limiting and high-pressure environment.

## Introduction

Ammonia oxidizing archaea (AOA) comprise one of the most successful archaeal phyla having colonized almost every imaginable oxic environment of the planet where they emerge as key players in the nitrogen cycle [1–6]. This includes the marine environment where they dominate archaeal communities associated with oxic sediments ranging from shallow estuaries to the open ocean [7–12], and from the surface layers all the way into the deep oceanic crust [13–15]. In these ecosystems they seem to play a critical role in the transformation of nitrogen compounds and control its partitioning into the bottom ocean and the underlying oceanic crust [12, 14–19]).

Studies from the North Atlantic and Pacific show that the composition of the sedimentary AOA population differs drastically from that in the overlying water suggesting distinct ecophysiological potential to colonise sedimentary environments, albeit all were found to belong to the order *Nitrosopumilales* (NP) [20]. Whereas the *amo*A-NP-gamma clade seems to be dominant and omnipresent in these oceans, irrespective of water depths, the *amo*A*-*NP-alpha clade represents the most abundant ecotypes in deep ocean waters [6, 7, 21–27] (nomenclature based *amo*A gene classification [28]). In contrast, dominant phylotypes in deep-sea sediments belong to the *amo*A-NP-theta and *amo*A-NP-delta clades [11, 12]. In addition, in cases of oligotrophic oceanic regions, these were detected throughout the sediment column and further into the underlying basaltic crust, even at depth where oxygen is below detection [7, 10–12, 14, 15]. In these, sites, they exhibit peaks of abundance and diversity at oxic/anoxic transition zones where increased energy availability is suggested to sustain the higher biomass of nitrifiers [12]. The abundance and distribution of *amo*A-NP-theta and -delta in the energy-starved subsurface suggest that they have adapted and evolved differently than their pelagic counterparts. These clades represent a yet unexplored diversity within *Nitrosopumilales*, and so far have no cultivated or genomic representative [28].

It has been suggested that the AOA common ancestor arose in terrestrial habitats (probably hot springs) where the AOA lineages diversified and then occupied different biomes (e.g. soils, hot springs and freshwater environments) before conquering estuarine and marine shallow water environments and finally, radiating into deeper waters as a result of the oxygenation of the deep ocean during the Neoproterozoic [29, 30] (Abby et al., submitted). AOA are generally well equipped for the manifold challenges of the oxic deep-sea surface and subsurface environment. They encode the most energy-efficient aerobic carbon fixation pathway [31] making them important primary producers in these environments [32, 33], and their high affinity for ammonia would enable them to utilize this scarce resource [34]. Nevertheless, deep pelagic as well as benthic AOA populations are reported to have the capability for mixotrophy as well, as indicated by uptake of labelled compounds and through the detection of uptake/assimilation genes for organic carbon and nitrogen compounds by shotgun metagenomics [10, 16, 23, 24, 33, 35–37]. Stimulation of autotrophic CO_2_ fixation by organic carbon was also shown by isotope labelling studies [32]. In the absence of genomic context however, virtually nothing of the above can be extrapolated to the metabolic potential or adaptations of ecotypes that dominate deep marine sediments, nor can their ecological boundaries be interpreted.

In this study, we address the question of what adaptations enabled specific AOA clades to inhabit bathyal and abyssal (i.e. deep-sea) marine sediments, and the significance of this in the context of thaumarchaeal evolution. To this end, we obtained the first high-quality metagenome-assembled genomes (MAGs) belonging to the so-far uncharacterized *amo*A-NP-theta and *amo*A-NP-delta clades from sediment cores obtained from the Mid-Atlantic Ridge flanks, and from the oligotrophic Pacific Ocean. We also describe two MAGs associated with a novel, deep-branching clade within the *Nitrosopumilales*, which we designate *amo*A-NP-iota (previously NP - *insertae sedis* [28]). The pivotal phylogenetic position of the latter and the distribution of all three clades in phylogenomic trees enables us to shed light on the evolutionary diversification of AOA into marine sediments, which seems much more complex than previously assumed and reveals unique, similar, and also overlapping adaptive strategies in all three clades.

## Materials and Methods

### Sampling of Atlantic and Pacific sediments

Oligotrophic sediment cores were retrieved from mid-ocean ridge flanks in the Atlantic Ocean: Hole U1383E (22^°^48.1’N, 46^°^03.2’W, 4425 m water depth) from North Pond by advanced piston coring during the International Ocean Drilling Program (IODP) Expedition 336 (2011), and GS14-GC08 (71°58.0’ N, 0°6.1’ E, 2476 m water depth) by gravity coring from the east flank of the central Mohns Ridge (2014) (Fig. 1a). Genomic DNA from four sediment horizons in each core were selected for metagenome sequencing, based on the published porewater geochemical data and 16S rRNA gene profiles [12, 38]. In particular, sediments of 0.1 m (oxic), 10.0 m (oxic), 22.0 m (oxic-anoxic transition zone; OATZ), and 29.5 m (anoxic-oxic transition zone; AOTZ) were selected from U1383E. Sediments of 0.1 meters below the seafloor (mbsf (oxic), 1.0 m (OATZ), 1.6 mbsf (nitrate-ammonium transition zone), and 2.5 mbsf (Mn-reduction zone) were selected from GS14-GC08. Detailed information about sampling sites, sampling procedure, 16S rRNA gene profiles and porewater analysis was published in [12, 38].

**Fig. 1.**
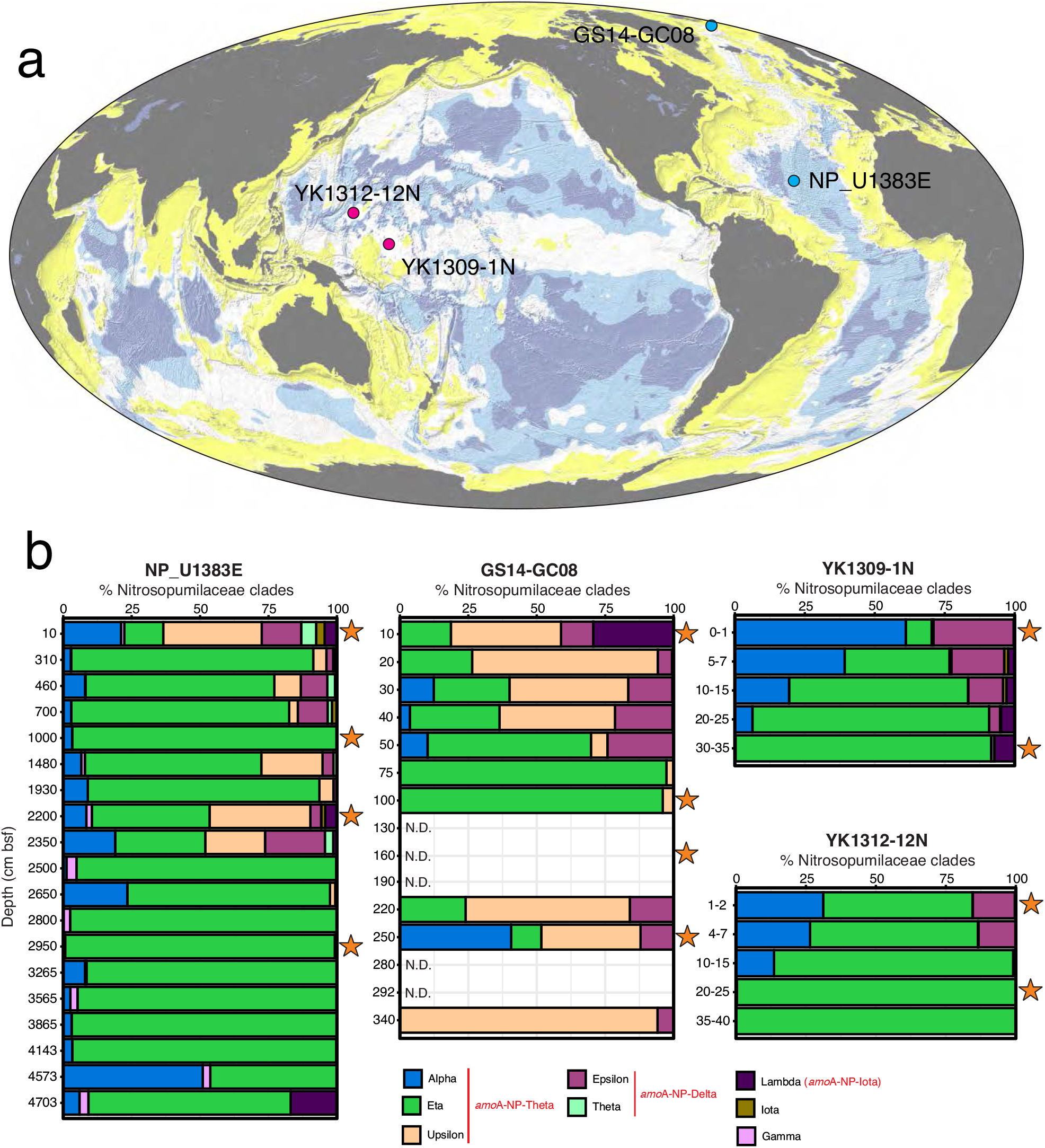
Study sites and the community structure of AOA. (**a**) Global bathymetric map showing the coring locations of the sediment cores used in this study, modified after [19]. The dark blue and light blue regions represent the minimum and maximum areas over which dissolved O_2_ is expected to penetrate throughout the sediment from seafloor to basement. (**b**) AOA community structure based on the 16S rRNA gene phylogeny. *Nitrosopumilaceae* 16S rRNA gene OTUs were classified based on their placements in the phylogenetic tree as described in [14]. In the figure key, the corresponding clades of the AOA *amo*A gene are shown in red. They are based on [55] and our phylogenetic analyses (unpublished data). The sediment horizons selected for metagenome sequencing are highlighted by orange stars. Data for NP_U1383E and GS14-GC08 were retrieved from [12].

Sediment cores (YK1309-1N and YK1312-12N) from the Pacific abyssal plain were collected using a push corer with a manned submersible *Shinkai6500* during the JAMSTEC cruises YK13-09 (September 2013: 01°15.0’N, 163°14.9’E, 4277 m water depth) and YK13-12 (November 2013: 11°59.9’N, 153°59.9’E, 5920 m water depth) of the *R/V Yokosuka,* respectively. Two sections from each core were selected for shotgun metagenomic sequencing: YK1309-1N-S000 (0-1 cmbsf), YK1309-1N-S300 (30-35 cmbsf), YK1312-12N-S010 (1-2 cmbsf) and YK1312-12N-S200 (20-25 cmbsf).

DNA of all samples was prepared using standard techniques and was sequenced on Illumina Hiseq2500, 16S rRNA gene amplicons were generated and sequenced using standard procedures (see Suppl. Material). Detailed information about sampling sites, sampling procedure, and geochemical analyses are also shown in Suppl. Material.

### Assembly and comparative genomics

All sequencing data were processed to remove illumina adapters and low quality reads using Trimmomatic [39] before *de novo* assembly using MEGAHIT [40] (k-mer length of 27-117). Binning of contigs of Pacific metagenomes was performed applying a contig dereplication and binning optimization tool [41] based on the binning output of CONCOCT [41], MetaBAT [42] and MaxBin2 [43] while contigs of Atlantic samples were binned with MaxBin2 [43] followed by a sequence of refinement steps for the Thaumarchaeal bins (see supplementary methods). Completeness and contamination of Pacific and Atlantic bins were evaluated with CheckM (“lineage_wf” parameter) [44]. Assemblies are available on NCBI (accession numbers pending).

A dataset consisting of 163,852 predicted proteins from 85 genomes (11 metagenome-assembled genomes (MAGs) reported here, 31 complete and near-complete AOA genomes and 43 MAGs or single-amplified genomes (SAGs) from NCBI or IMG) was collected for this study (Table S1). Annotation of the MAGs assembled in this study was performed automatically using the Microscope annotation platform from Genoscope [45], followed by extensive manual curation. NCBI annotations were supplemented with arCOG assignments from the archaeal Clusters of Orthologous Genes database (2018 release) [46] using COGsoft [47] (e-value of 10^−10^). We clustered the protein dataset into protein families based on sequence identity (35 %) and alignment coverage (70 %) using CD-Hit V4.8.1 [48] (“-c 0.35 -aL 0.7 -aS 0.7”) (Table S3).

### Selection of markers and phylogenomic tree

The identification of markers to perform the phylogenomic tree reconstruction was based on the phylogenomic workflow proposed by [49] (E-value 10^−10^) using the archaeal single-copy gene collection [50]. We selected 79 markers (Table S2), present in at least 70 of the 85 genomes used in this study. Each protein family was aligned using MAFFT v7 (“--maxiterate 1000–localpair”) [51] and trimmed with BMGE [52]. The concatenated alignment was used to reconstruct a Maximum likelihood phylogenomic tree in IQTREE (v2.0-rc1) [53] under the LG+C20+F+G model with 1,000 ultrafast bootstrap replicates. For *amo*A phylogeny and detailed methodological procedures, see Supplementary Material.

## Results and Discussion

### Distribution of AOA in deep marine sediments

We examined the overall community structure of AOA (all affiliated to the family *Ca.* Nitrosopumilaceae) in these sediments by analyzing 16S rRNA gene amplicon sequencing data generated in this study for the Pacific cores and previously described for the Atlantic cores [12]. AOA communities in sediment horizons deeper than 10 cm were all dominated by the so-called 16S-NP-eta and/or 16S-NP-upsilon and 16S-NP-alpha clades [55], which together correspond to the *amo*A-NP-theta clade (Fig. 1b) [28]. In addition, AOA affiliated to the 16S-NP-epsilon clade (corresponding to the *amo*A-NP-delta clade, Fig 2a) were also repeatedly detected with percentages <25% in the upper portions of these cores (Fig. 1b). Finally, the 16S*-*NP-lambda clade (now renamed to *amo*A-NP-iota, see below) was also detected as a minor clade in all cores except YK1312-12N, but was notably abundant in the uppermost horizon of GS14-GC08 (29% of the total AOA community, Fig. 1b).

**Figure 2.**
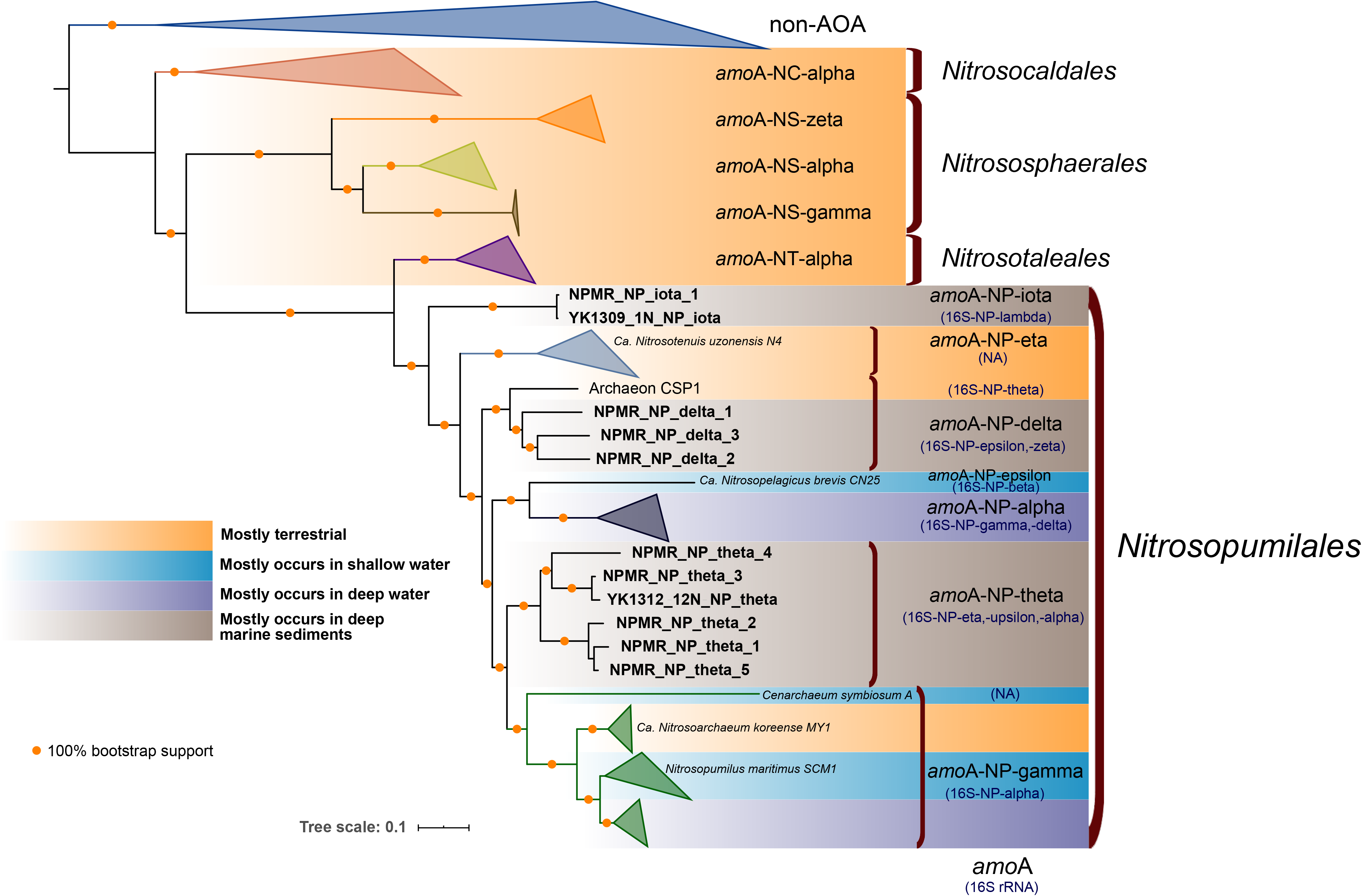
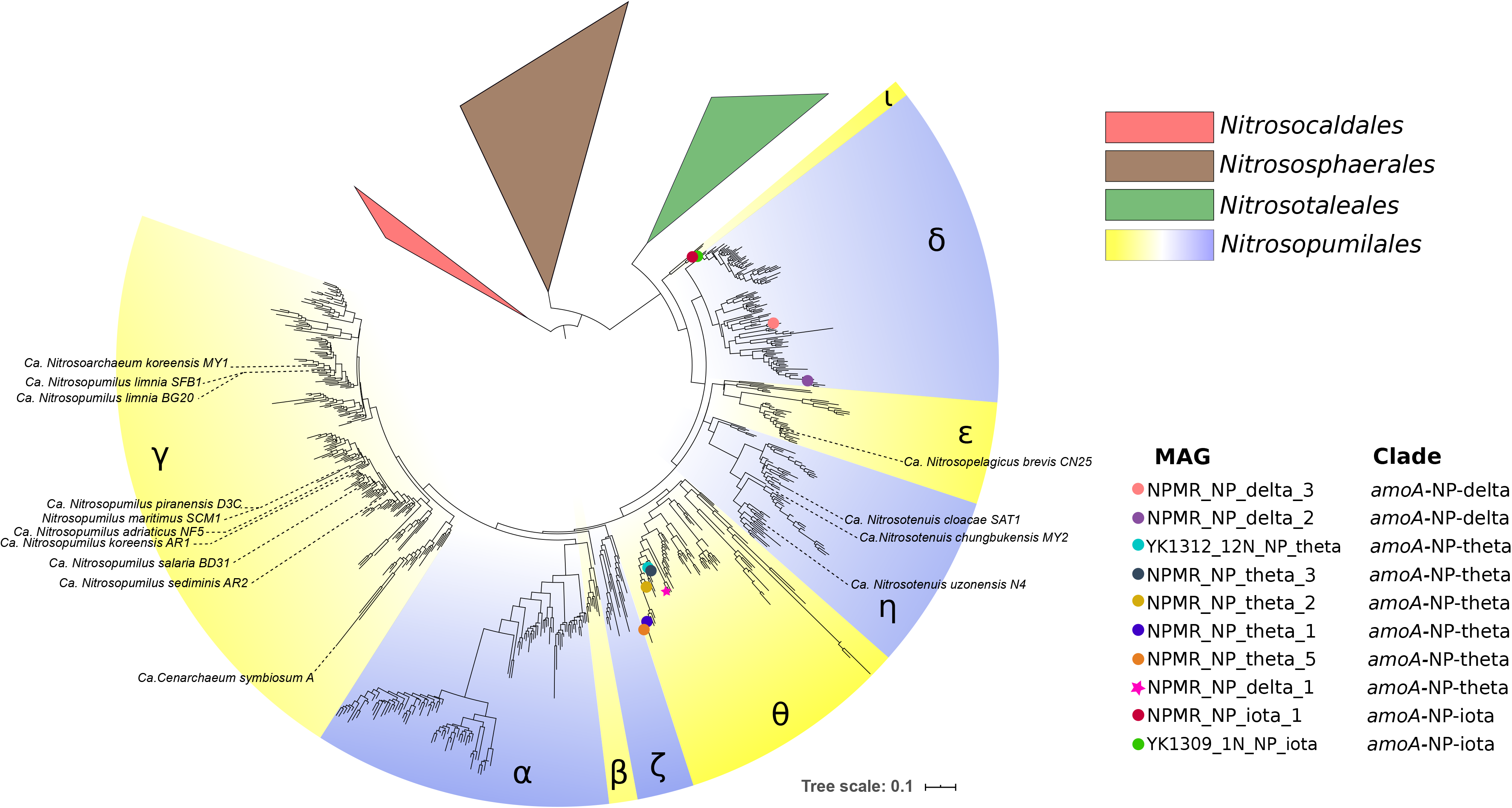
**a) Phylogenomic tree of AOA and non-AOA Thaumarchaeota.** The phylogenomic tree was reconstructed based on the concatenated alignment of 79 markers comprising 7,485 amino acid sites using a maximum likelihood approach (see Materials and methods). Yellow circles represent 100 % bootstrap support of nodes. The metagenome-assembled genomes (MAGs) reported in this study are shown in bold. Representative genomes of NP-subclades were added for subclade clarification. Information about the ecological distribution of AOA clades is provided. The scale bar indicates the number of substitutions per amino acid site. In most cases congruence between *amo*A-based and 16S rRNA-based clades was inferred from complete genomes or MAGs where both genes were present [14, 22, 23]. For other cases: 1) *amo*A-NP-iota/16S-NP-lambda pair: the congruence of these two clades is inferred from phylogenetic trees and congruence in relative abundances (Zhao and Jorgensen, unpublished data) of sediment layers with high and changing relative abundances; 2) *amo*A*-*NP-delta/16S-NP-epsilon,-zeta pair: The congruence of these two clades is based on the MAG of CSP1 (with both genes), the close relationship of 16S-NP-epsilon with CSP1 and the occurrence pattern of this clade in marine sediments (see [12], and data not shown); and 3) *amo*A-NP-theta/16S-NP-alpha,-eta,-upsilon pair: congruence based on [14], own phylogenetic reconstructions (unpublished data) and relative abundances in marine sediment layers. Abbreviations: NA; not assigned. **b) Phylogeny of** *amo***A sequences.** The ML phylogenetic tree was reconstructed using 584 nucleotide sites. Colored circles represent MAGs from this study. The star symbol represents the NPMR_NP_delta_1 bin. The incongruity of its phylogenetic placement based on the *amo*A gene and the phylogenomic analysis of concatenated markers is discussed in the text. Greek letters represent the *amo*A-based annotation of AOA subclades as in [28]. The scale bar indicates the number of substitutions per nucleotide site.

### Phylogenomic analysis and taxonomic placement reveal MAGs from three independent *amo*A clades

We obtained a total of 11 AOA metagenome-assembled genomes (MAGs), 9 from Atlantic and 2 from Pacific sediment samples (sequenced horizons are marked by stars in Fig. 1b). Despite high sequencing depths and high AOA abundance in the sample, based on 16S rRNA gene reads in the metagenomes (10 - 16 % in the Pacific cores and 6.8-18.8% in the Atlantic cores), generation of good quality bins was extremely challenging, possibly due to high microdiversity (41 OTUs) within *Ca.* Nitrosopumilales as observed earlier [12]. Eventually we obtained 4 MAGs with >90 % completeness and 3 MAGs with >80 % completeness, all with contamination levels ≤ 5%, which we consider high quality MAGs in this study, as well as four additional medium quality MAGs (66 to 76% completeness, up to 6.3% contamination level) (Table1). The MAGs genome sizes (0.61 to 1.52 Mb) and GC contents (34.66 ± 0.72%) are in accordance with previous reports of free-living *Ca.* Nitrosopumilales [56].

**Table 1.** Statistics of deep marine sediments-derived MAGs. High-quality MAGs (> 80 % completeness and ~ 5% contamination or below) are marked with an asterisk symbol.

| Genome Bin | AOA clade | Depth (m) | Depth (m<br>below sea<br>floor) | Contigs (No.) | Genome size<br>(bp) | Proteins<br>(No.) | Longest<br>contig (bp) | N50 | GC (%) | Completeness<br>(%) | Contamination<br>(%) | Presence/<br>absence of 16S<br>rRNA gene |
| --- | --- | --- | --- | --- | --- | --- | --- | --- | --- | --- | --- | --- |
| NPMR_NP_delta_1* | NP-delta | 4,425 | 0.1 | 83 | 1,173,997 | 1,467 | 79,876 | 28,788 | 34.00 | 86.23 | 5.42 | - |
| NPMR_NP_delta_2 | NP-delta | 4,425 | 0.1 | 270 | 610,368 | 870 | 12,327 | 3,054 | 33.87 | 67.88 | 0.71 | - |
| NPMR_NP_delta_3 | NP-delta | 4,425 | 0.1 | 143 | 937,620 | 1,222 | 58,541 | 15,484 | 34.22 | 66.29 | 3.4 | - |
| NPMR_NP_theta_1* | NP-theta | 4,425 | 1 | 176 | 1,524,500 | 2,000 | 40,454 | 14,847 | 35.06 | 99.51 | 2.91 | - |
| NPMR_NP_theta_2* | NP-theta | 4,425 | 22 | 298 | 1,201,755 | 1,671 | 26,499 | 5,123 | 34.98 | 93.69 | 4.05 | - |
| YK1312_12N_NP_theta* | NP-theta | 5,920 | 0.2 | 299 | 1,221,321 | 1,695 | 20,455 | 5,469 | 34.08 | 88.83 | 5.34 | - |
| NPMR_NP_theta_3* | NP-theta | 2,476 | 0.1 | 552 | 1,069,514 | 1,706 | 14,642 | 2,738 | 34.37 | 83.58 | 3.87 | + |
| NPMR_NP_theta_4 | NP-theta | 2,476 | 0.1 | 232 | 1,040,901 | 1,408 | 21,324 | 6,332 | 33.98 | 76.24 | 6.31 | - |
| NPMR_NP_theta_5 | NP-theta | 4,425 | 0.1 | 274 | 697,980 | 970 | 14,641 | 3,916 | 35.14 | 75.01 | 1.21 | - |
| NPMR_NP_iota_1* | NP-iota | 2,476 | 0.1 | 68 | 1,121,898 | 1,402 | 75,887 | 30,338 | 35.83 | 98.54 | 0.97 | - |
| YK1309_1N_NP_iota* | NP-iota | 4,277 | 0.3 | 227 | 1,298,549 | 1,700 | 40,725 | 9,447 | 35.76 | 96.6 | 3.24 | - |

In order to study the evolution of AOA and place our deep marine sediments-derived MAGs in a phylogenetic context, we reconstructed a maximum-likelihood (ML) phylogenomic tree (Fig. 2a) using 79 concatenated single-copy markers from our entire dataset of 85 complete genomes, MAGs and SAGs representing a broad diversity of habitats (Table S1).

In addition, we performed an *amo*A-based phylogeny as in [28] in order to assign a taxonomical rank and a respective AOA clade to our MAGs (Fig. 2b). Both trees showed similar clustering of MAGs into *Ca.* Nitrosopumilales subclades except for NPMR_NP_delta_1 (see discussion in Supplementary Information) which based on the *amo*A tree clustered within the *amo*A-NP-theta clade but the more robust phylogenomic analysis strongly suggests that it belongs to the *amo*A-NP-delta subclade. The only 16S rRNA gene recovered in MAG NPMR_NP_theta_3 is affiliated with a subclade of 16S-NP-alpha exclusively found in marine sediment (not shown).

Our phylogenomic tree revealed that the 11 AOA MAGs reported here represent the dominant AOA observed in our study (Fig. 1b) and form three well supported monophyletic groups, of which no cultured representative has been reported yet (Fig. 2a). Six MAGs represent the first genomic assemblies from the *amo*A-NP-theta lineage, one of the most dominant AOA groups in marine sediments and also found to be abundant in the crust below [14, 28]. Three MAGs are affiliated to *amo*A-NP-delta, the second most abundant AOA clade in marine sediments, and are the first marine sediment representatives of this clade, which includes a single other MAG (Archaeon CSP1) assembled from river aquifer sediments [57].

Two bins (NPMR_NP_iota_1 and YK1309_NP_iota) clustered together forming a third sediment-dwelling clade, sister to all *Ca.* Nitrosopumilales, which earlier escaped taxonomic assignment as it was only identified based on singular *amo*A sequences and hence had been designated *incertae sedis* [28]. Pairwise average nucleotide identity (ANI) comparisons (Fig. S2), indicate that these two bins share > 70 % ANI with the other NP-MAGs recovered in this study (*amo*A-NP-delta and *amo*A-NP-theta MAGs sharing 73 - 79 % ANI). A comparison of conserved protein families among all NP subclades indicated that this group harbors 320 out of 336 protein families that seem to be part of the *Ca.* Nitrosopumilales core proteome, as opposed to only 260 of these protein families being present in the *Ca.* Nitrosotaleales, the sister lineage to all NP (Fig. 3). Moreover, environmental *amo*A sequences suggested that this clade might be restricted to deep-sea sediments, an ecological specialization only found in *Ca.* Nitrosopumilales. Taken together, this early-branching clade seems to be a new NP subclade and we propose the designation *amo*A*-NP-*iota (as it forms the ninth NP-clade following the taxonomy of [28]).

**Figure 3.**
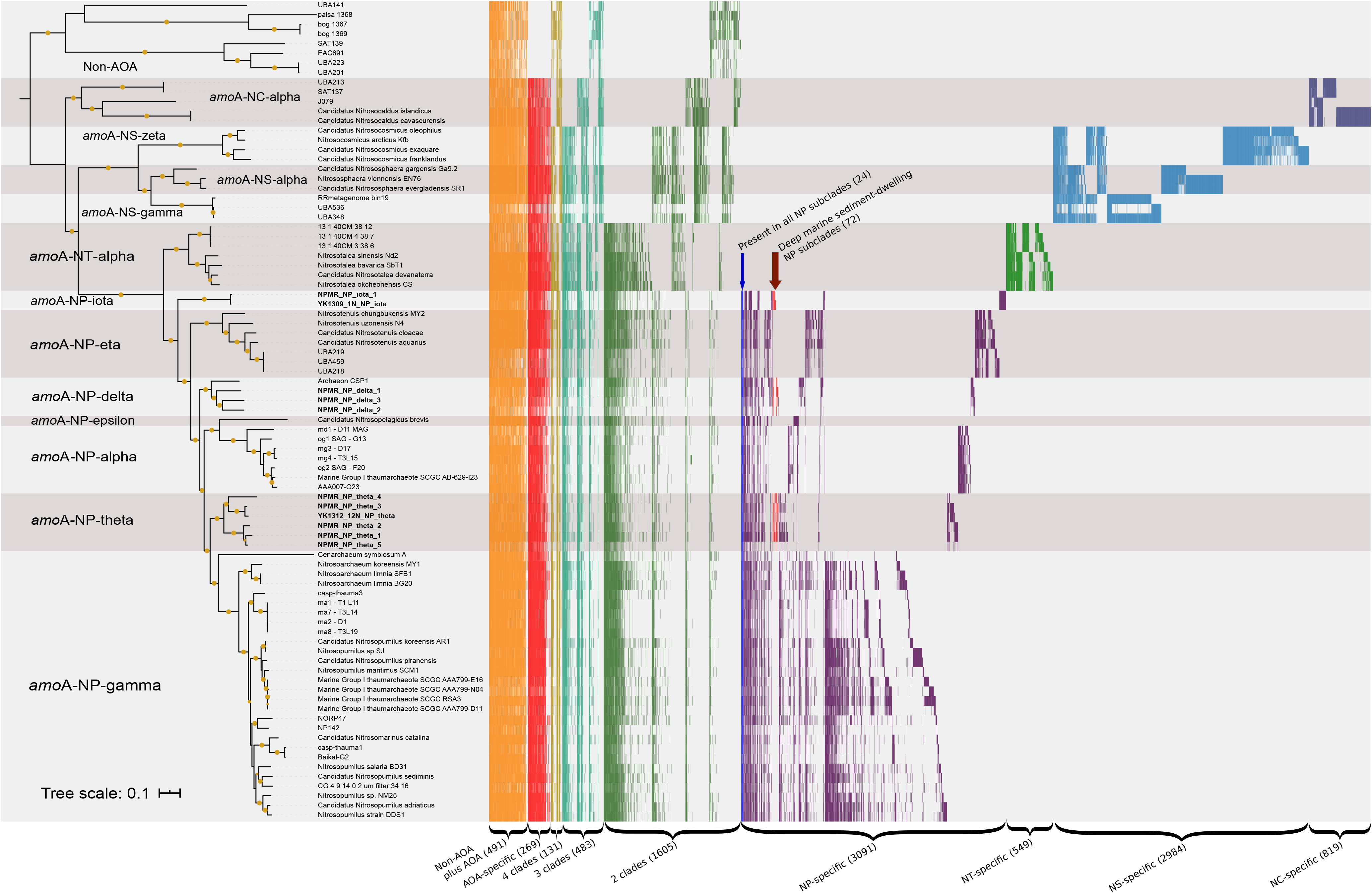
Comparative analysis of the presence/absence of protein clusters among AOA. Each bar (x axis) represents that the putative protein family is encoded in the genome (y axis). 10, 422 clusters with at least two different genomes are depicted. Non-AOA-specific clusters were excluded from the visualization. Yellow circles represent > 95 % bootstrap support of nodes. Clusters are ordered based on their distribution pattern from the most widespread to the most uncommon: first, across all lineages and then intra-lineage. Groups of clusters have different color codes for better visualization. NP represents Nitrosopumilales; NC, Nitrosocaldades; NS, Nitrososphaerales; NT, Nitrosotaleales.

### Three independent radiations of AOA into marine sediments

While MAGs and SAGs of AOA from bathypelagic (1000-4000m) to abyssopelagic (4000-6000m) and hadopelagic (6000-11000m) environments have been reported previously [23, 24, 58] and shotgun metagenomic analyses of deep-sea sediments have been performed [10, 59, 60], the MAGs reported in this study represent to our knowledge the first high-quality AOA genomes from bathyal and abyssal sediments (> 2000 m depth). Together with our taxon-enriched phylogenomic analyses they shed new light on the ecological transitions and niche differentiation undergone by AOA.

While our phylogenomic tree supports that AOA likely appeared in terrestrial habitats first (Fig. 2a), the sequence and number of colonization events of marine environments seem to be more complex than previously proposed [29]. Importantly, our results suggest that both the deep-water adapted AOA as well as the deep-sediment adapted AOA are polyphyletic. The colonization of deep waters, i.e. pelagic organisms, might have occurred independently at least twice in the evolution of the *Ca*. Nitrosopumilales, once at the origin of *amo*A-NP-alpha clade and the other during the diversification of *amo*A-NP-gamma (Fig. 2a). Interestingly, the *amo*A-NP-gamma clade which is one of the most diverse *Ca*. Nitrosopumilales subclades [28] has undergone particular habitat transitions and niche occupation [28]. Distinct shallow water *amo*A-NP-gamma species have established independently symbiotic associations with sponges [56, 61] while the sub-lineage leading to the soil isolate *Ca*. Nitrosarchaeum koreense [62] could have evolved from an estuarine or shallow water ancestor suggesting a recolonization of land (Fig. 2a).

Regarding the origin of deep sediments-dwelling AOA, the *amo*A-NP-theta lineage branches within mostly marine NP clades (Fig. 2a), suggesting that this lineage might have evolved from a pelagic marine ancestor. However, there is no evolutionary link between any pelagic AOA and the NP-delta clade, as the latter have been mostly retrieved in estuarine and deep marine sediments [57]. The most parsimonious evolutionary scenario would be that this group underwent a direct transition from estuarine sediments to marine sediments during its diversification.

Similarly, the newly proposed clade NP-iota, the earliest branching Nitrosopumilales which has so far exclusively been detected in marine sediments [28] does not seem to be closely related to pelagic Nitrosopumilales but emerges instead among terrestrial clades (*Ca*. Nitrosotaleales and NP-Eta). Although, it is possible, that pelagic AOA closely related to *amo*A-NP-iota may be detected in further environmental surveys or that respective pelagic lineages got extinct, the *amo*A-NP-iota clade might as well have developed from terrestrial-estuarine organisms, as discussed for *amo*A-NP-delta above.

### Comparative genomics of deep-sea sediment AOA

We constructed a total of 33,442 protein families from our taxon-enriched genome dataset representing a wide variety of ecological environments (see Materials and Methods and Table S1). From these, 12,137 have representatives from at least two different genomes. In our analysis, the AOA core proteome comprises 760 protein families present in at least one genome of each of the four major AOA lineages: *Ca*. Nitrosocaldales, *Nitrososphaerales*, *Ca.* Nitrosotaleales and *Ca.* Nitrosopumilales (Fig. 3, Table S3). Thus, our results are similar to previous estimations of the AOA core genome (743 gene families) [63], and slightly lower than our own earlier estimate of 860 gene families (based on only 7 genomes [64]). Only 269 of the core AOA families seem to be AOA-specific (Fig. 3, Table S3). Only 123 out of these 269 families were found to be present in >50 % of the genomes in each of the four AOA orders (a relatively low threshold to account for the incompleteness of MAGs), suggesting a relatively low degree of conservation within these lineages. These results imply great intra-order genomic variability and important differential gene loss among subclades and across genomes during the evolution and diversification of AOA. For instance, despite the fact that Nitrosopumilales have 3091 specific families with proteins encoded in at least two genomes and present in one or more NP-subclades, a subset of solely 24 families were conserved in all 7 NP-subclades (Fig. 3). Considering the very relaxed criteria used, this is an astonishingly small number of conserved families in all 7 NP-subclades.

To identify possible specific adaptations of AOA to deep marine sediments, we searched for families present in at least two of the three marine sediments clades represented by our 11 MAGs (i.e. *amo*A-NP-theta, -delta and -iota), to the exclusion of all the other genomes analyzed in this study (Fig S2, Table S1). 72 families were identified (Fig. 3), of which only 25 % (18 families) could be functionally annotated (Table S3) and were classified into the following categories: information processing systems (7), metabolism (5) and cellular processes (6). Some of these 18 families had functional equivalents in most if not all AOA (e.g. RadA homologs). We additionally found 41 families shared predominantly between NP sub-clades with deep ocean (>1000m) representatives (i.e. *amo*A-NP-alpha, NP-gamma sublineages recovered from the Mariana, Izu-Ogasawara Trenches and the Red Sea [23, 58], NP-theta, NP-iota and NP-delta), to the exclusion of all other NP sub-clades. From these 41 families, 18 have functional annotation: information processing systems (8), metabolism (4) and cellular processes (6). Families with functional significance specific to marine sediments, such as a putative lactate racemase, or those shared with deep ocean MAGs, are discussed below. Families identified in deep ocean MAGs but not found in the sediment clades are still depicted in Fig. S4 for comparative purposes. Additionally, we investigated the number of clusters shared between deep sediments-derived MAGs and the terrestrial (present in soils and sediments) lineage *Nitrososphaerales*, to the exclusion of all other AOA lineages and NP-subclades. Interestingly they share only one protein family related to coenzyme F_420_-dependent luciferase-like oxidoreductases.

### Metabolic reconstruction of the *amo*A-NP-theta, *amo*A-NP-delta, *amo*A-NP-iota clades

The full annotations for all genes and pathways discussed in the following section can be found in Table S4.

### Central energy and carbon metabolism

All three sediment clades (i.e. *amo*A-NP-delta, *amo*A-NP-theta, *amo*A-NP-iota) encode complete sets of genes involved in ammonia oxidation, namely *amo*AXCB in the typical organization observed in other *Nitrosopumilales* (Fig. 5, Table S3) [21, 65, 66]. Missing subunits in certain MAGs seem to be due to genome incompleteness. A nitrite reductase (NirK) homolog is present, as well as multiple blue copper domain proteins putatively functioning as electron carriers. All clades encode a single high-affinity ammonia transporter family protein (Amt), as opposed to two Amt transporters of differing affinities found in other AOA. This could represent an adaptation to an oligotrophic environment [67]. Four out of six NP-theta MAGs and all *amo*A-NP-delta MAGs encode complete or near-complete urease operons (Fig. 4, Table S3). Together with a putative nitrilase (Nit1, conserved in AOA) and a putative omega-amidase (Nit2, present in *amo*A-NP-theta, -delta, -eta, -gamma), these genes indicate expanded substrate utilization capabilities for ammonia (and CO_2_) generation by cleaving urea, nitriles and dicarboxylic acid monoamides. Utilization of organic nitrogen compounds is a feature shared with other NP-clades that include deep-sea lineages and previously described for subseafloor AOA (Fig. 4, 5, S4) [10, 23, 24, 33].

**Figure 4.**
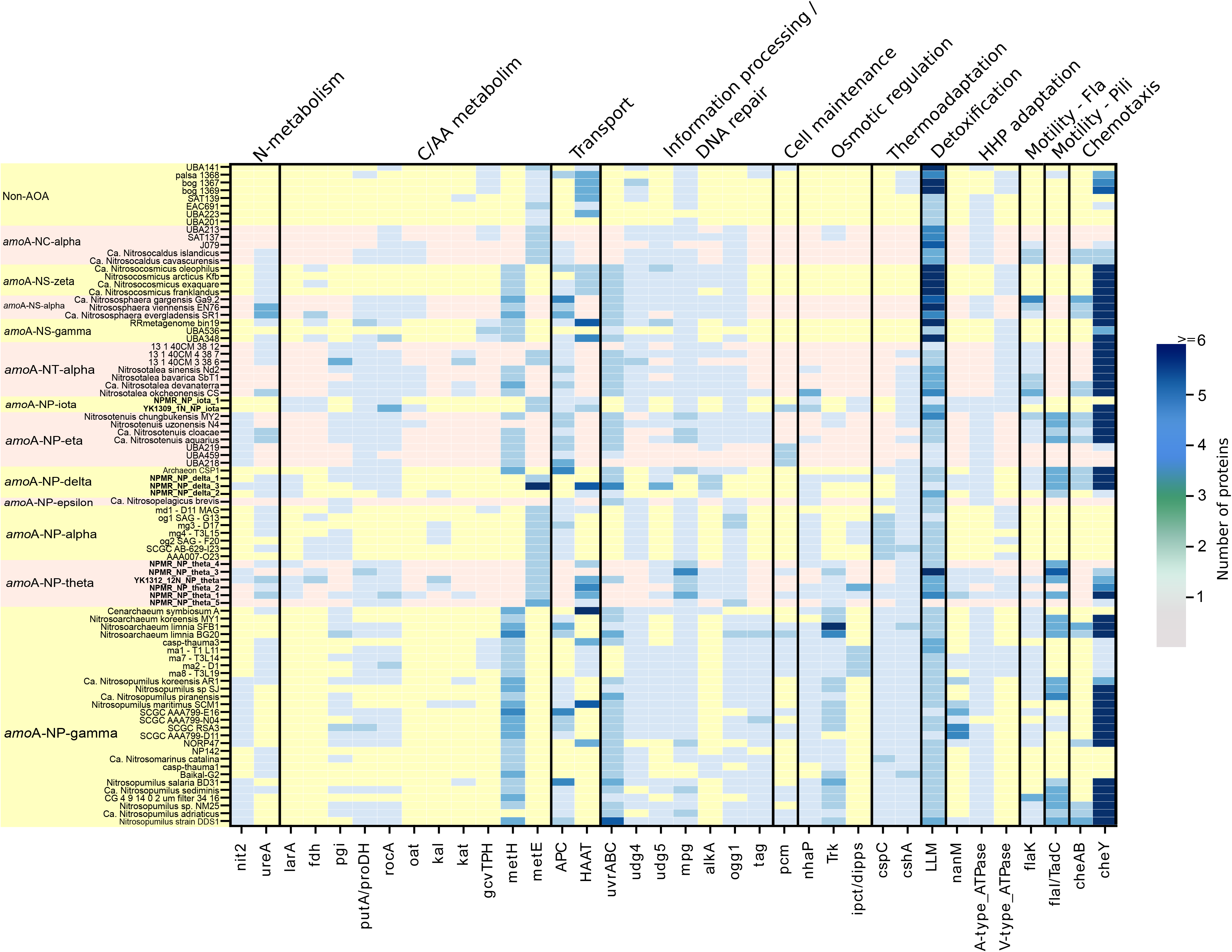
Heatmap depicting the distribution and abundance of genes involved in the main functional categories discussed in the text. Abbreviations: nit2, nitrilase/omega-amidase; ureA, urease subunit gamma; fdh, formate dehydrogenase; larA, lactate racemase; pgi, phosphoglucose isomerase; proDH, proline dehydrogenase; rocA, 1-pyrroline-5-carboxylate dehydrogenase; oat, putative ornithine--oxo-glutarate aminotransferase/class III aminotransferase; kal, 3-aminobutyryl-CoA ammonia lyase; kat, putative 3-aminobutyryl-CoA aminotransferase; gvtTPH, glycine cleavage system proteins T/P/H; metH, methionine synthase II (cobalamin-independent); metE, methionine synthase I (cobalamin-dependent); APC, Amino Acid-Polyamine-Organocation Transporter Family; HAAT, the Hydrophobic Amino Acid Uptake Transporter (HAAT) Family; uvrABC, the Uvr excision repair system endonucleases ABC; udg4/5, Uracil DNA glycosylase family 4/5; mpg, methylpurine/alkyladenine-DNA glycosylase; ogg1, 8-oxoguanine DNA glycosylase; alkA, DNA-3-methyladenine glycosylase; tag, 3-methyladenine DNA glycosylase; pcm, protein-L-isoaspartate carboxylmethyltransferase; nhaP, the Monovalent Cation:Proton Antiporter-1 (CPA1) Family; Trk, the K+ Transporter (Trk) Family; ipct/dipps, bifunctional CTP:inositol-1-phosphate cytidylyltransferase/di-myo-inositol-1,3’-phosphate-1’-phosphate synthase; cspC, cold-shock protein A; cshA, cold-shock DEAD-box protein A; LLM, luciferase-like monooxygenase family protein; nanM, N-acetylneuraminic acid mutarotase; flaK, archaeal preflagellin peptidase FlaK; cheY, chemotaxis response regulator CheY; cheAB, chemotactic sensor histidine kinase cheA & methylesterase cheB; All locus tags and cluster information are in Supplementary tables 3 & 4. An extended version of the heatmap is in Fig S3

**Figure 5.**
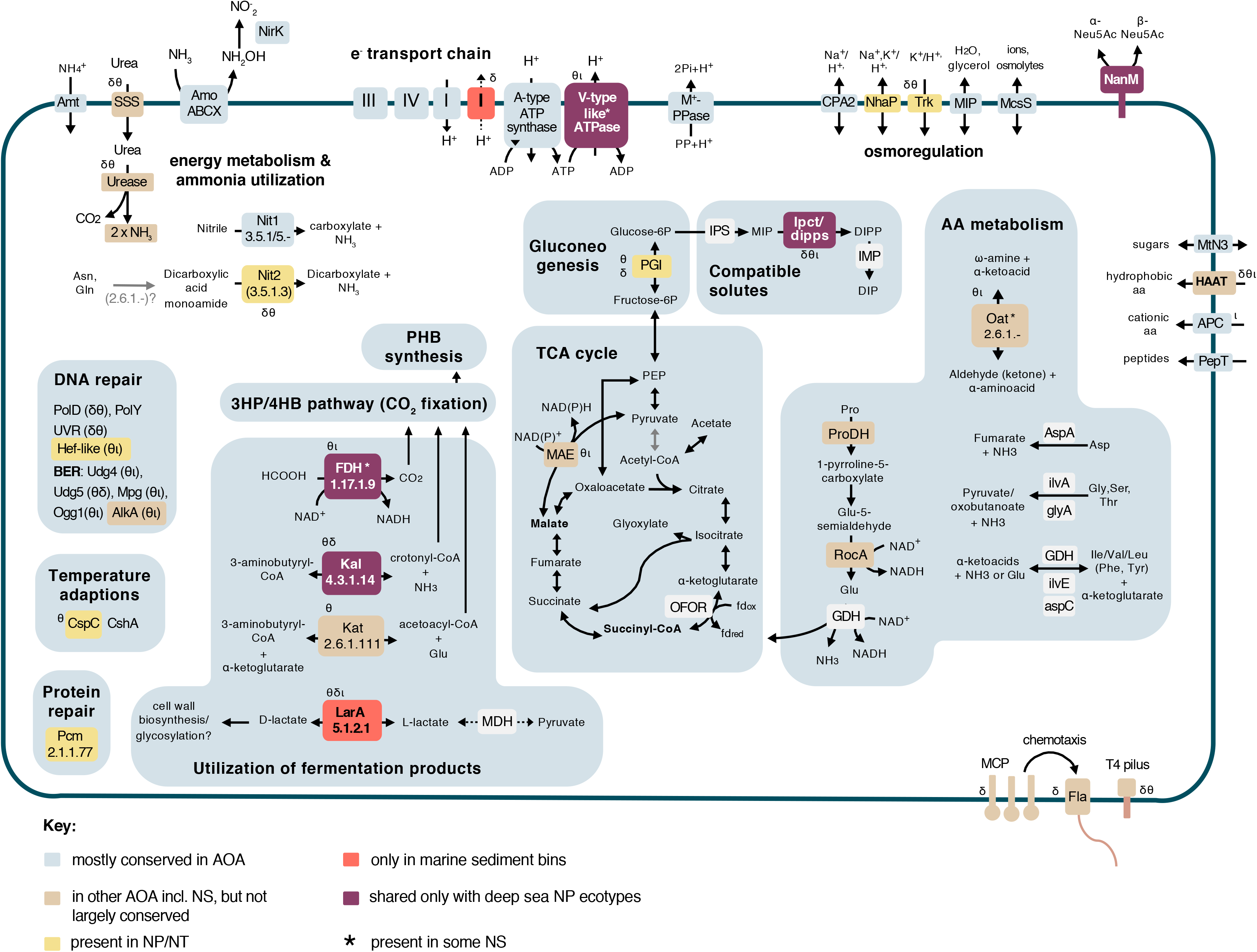
Metabolic reconstruction of *amo*A-NP-theta, *amo*A-NP-delta and *amo*A-NP-iota AOA. Schematic reconstruction of the predicted metabolic modules in the sediment MAGs, as discussed in the text. Color code of enzymes/complexes indicates conservation status in AOA. Unless specified by greek letters (θδι), enzymes/modules are present in all sediment clades. Dashed lines indicate hypothetical reactions. Gray arrow indicates an alternative OFOR reaction. Complexes of the electron transport chain are labelled with roman numerals. Transporters are named according to TCDB classification. Enzymes, gene accession numbers and transporter classes are also listed in Supplementary Table 3. Abbreviations: Amo, ammonia monooxygenase; NirK, nitrite reductase; Nit1, nitrilase; Nit2, nitrilase/omega-amidase; AA, amino acid; Fdh, formate dehydrogenase; Kal, 3-aminobutyryl-CoA ammonia lyase; Kat, putative 3-aminobutyryl-CoA aminotransferase; Lar, lactate racemase; Pcm, protein-L-isoaspartate carboxylmethyltransferase; MCP, methyl-accepting chemotaxis protein; Fla, archaellum; PolD, polymerase family D; PolY, translesion polymerase family Y; UVR, excision repair system; Hef-like, Hef/FANCM/Mph1-like helicase; BER, base-excision repair; Udg4/5, Uracil DNA glycosylase family 4/5; Mpg, methylpurine/alkyladenine-DNA glycosylase; Ogg1, 8-oxoguanine DNA glycosylase; AlkA, DNA-3-methyladenine glycosylase; CspA, cold-shock protein A; CshA, cold-shock DEAD-box protein A; Pcm, protein-L-isoaspartate carboxylmethyltransferase; PHB, polyhydroxybutyrate; MAE, malic enzyme; OFOR, 2-oxoacid:ferredoxin oxidoreductase; PGI, phosphoglucose isomerase; IPS, myo-inositol-1-phosphate synthase; ipct/dipps, bifunctional CTP:inositol-1-phosphate cytidylyltransferase/di-myo-inositol-1,3’-phosphate-1’-phosphate synthase; IMP, DIPP phosphatase; ProDH, proline dehydrogenase; RocA, 1-pyrroline-5-carboxylate dehydrogenase; glutamate dehydrogenase; oat, putative ornithine--oxo-glutarate aminotransferase/class III aminotransferase; aspA, aspartate ammonia-lyase; ilvA, threonine/serine ammonia-lyase; glyA, serine/glycine hydroxymethyltransferase; ilvE, branched-chain-amino-acid transaminase; aspC, aspartate/tyrosine/aromatic aminotransferase; MCO1, multicopper oxidase family 1. NanM, N-acetylneuraminic acid mutarotase; Transporters are named according to TCDB classification (supplementary table 4).

All three sediment clades encode full gene sets for electron transfer to O_2_ via NADH dehydrogenase (complex I), type *bc*_1_ complex III, and a heme-copper terminal oxidase (complex IV) (Fig. 5, Table S3). No alternative complexes using a different electron acceptor were identified.

All three sediment clades encode the full repertoire conserved among AOA for autotrophic carbon fixation via the 3-hydroxypropionate/4-hydroxybutyrate (3HP/4HB) cycle and carbon metabolism through oxidative TCA and gluconeogenesis up to the formation of glucose-6P via a phosphoglucose isomerase (not present in NS) homolog in NP-theta, NP-delta (Fig. 5, Table S3) [31, 65, 68]. A malic enzyme, enabling the formation of pyruvate from malate with the concomitant generation of NAD(P)H expands metabolic capacities in NP-theta and NP-iota (also in some other AOA, Fig. 5). As most AOA, all sediment clades have the capacity to synthesize polyhydroxybutyrate (PHB) storage compounds, an obvious advantage in an oligotrophic environment [69].

Complete or near-complete amino acid biosynthesis pathways as well as vitamins (including vitamin B12) are present in all three sediment clades, as in other AOA (Table S3). As observed in NP-alpha representatives [23], the sediment clades use the B12-independent pathway for methionine biosynthesis (*metE*) (Fig. 4). Albeit this being a less catalytically efficient enzyme than the B12-dependent *metH* present in all other AOA, it is nevertheless much less costly energetically [70], and would therefore be an advantage in an energy-limiting environment where maintenance rather than fast growth is the norm [71].

### Utilization of exogenous organic compounds

All three sediment lineages seem to be capable of utilizing exogenous organic compounds from fermentation processes such as formate, lactate and 3-aminobutyryl-CoA as a source of carbon, nitrogen and reductive potential. This finding expands the range of organic carbon and nitrogen substrates suggested earlier for deep ocean AOA (previously comprising amino-acids, peptides and compatible solutes) and reinforces their role as key players in nutrient cycling in these biomes [6, 10, 23, 24, 27, 33].

A putative soluble NAD^+^-dependent formate dehydrogenase (Fdh), distinct from the iron-sulfur/molybdenum containing Fdh enzymes traditionally found as part of formate-hydrogen lyase systems [72], is found in the NP-theta and NP-iota clades (as well as in certain NS representatives and NP-alpha Fig. 4, 5). However, no additional hydrogenases or known formate transport systems were identified in the marine sediment bins. Our phylogenetic analysis (Fig. S3) indicates that the enzyme is a bona fide NAD^+^-dependent Fdh within the superfamily of D-2-hydroxyacid dehydrogenases [73]. This indicates the capacity to use formate for supplementing CO_2_ needs while concomitantly supplying reducing equivalents (as in methylotrophs [74, 75]).

A putative 3-aminobutyryl-CoA aminotransferase (Kat, EC 2.6.1.111) and a 3-aminobutyryl-CoA ammonia lyase (Kal, EC 4.3.1.14) were identified in the NP-theta and NP-delta bins, and are also found in some NP-alpha, NP-gamma and NT lineages (Fig. 4, 5). These enzymes participate in lysine fermentation pathway variants in fermentative bacteria [76]. Although the key pathway enzymes are not present in the sediment bins or any other AOA, this intermediate compound (3-aminobutyryl-CoA) could be scavenged from fermenting microorganisms in the sediment community. Both enzymes can remove ammonia from 3-aminobutyryl-CoA either by transferring it to α-ketoglutarate resulting in the formation of acetoacyl-CoA and glutamate (Kat), or by an elimination reaction that produces crotonyl-CoA and free ammonia (Kal). Both products are intermediates of the 3HB/4HP (CO_2_ fixation-) pathway and could be processed accordingly, generating reducing potential in the subsequent steps.

The presence of a putative lactate racemase family protein (LarA), specific to the NP-theta, delta and iota clades (Fig. 4, 5) suggests that lactate is another fermentation product that could be utilized by these lineages. This is one of the very few protein families with a putative function prediction shared specifically between the sediment AOA clades to the exclusion of all other AOA, suggesting an essential role. LarA in lactobacilli catalyzes the interconversion of D- and L-lactate, ensuring an adequate supply of D-lactate which is an important cell wall component conferring resistance to vancomycin [77] (see Supplementary Information). Given the importance of cell envelope maintenance in the adverse conditions of the sediments, it is possible that D-lactate has a similar use in sediment AOA, conferring resistance to exogenous toxic compounds. Alternatively, the lactate dehydrogenase-like malate dehydrogenase homologs found in AOA possess features indicating that they could have a broad substrate specificity, being able to utilize pyruvate in addition to oxaloacetate, and producing the L-stereoisomers of the products (see Supplementary Information and Fig. S5), with the concomitant reduction of NAD^+^ [78].

As mentioned above, the only protein family specifically shared among the sediment MAGs and the terrestrial NS lineages, is an F_420_-dependent luciferase-like oxidoreductase. While the metabolic role of these proteins in AOA in general is still unclear, the ability to degrade recalcitrant carbon via oxygenases in a manner similar to terrestrial organisms has been observed in sediment communities [60, 69], and is proposed to provide an opportunistic advantage for expanded substrate utilization in limiting conditions.

### Adaptations to low energy and high pressure environments

Deep-sea sedimentary environments found under the oligotrophic ocean present manifold challenges to microbial life, namely energy limitation, high hydrostatic pressure (HHP), low temperatures (< 4^°^C) and potential microoxic or anoxic conditions detrimental to aerobic metabolisms [18, 19, 71, 81–84]. Microorganisms respond with global metabolic changes rather than stress responses [85], some of which are found in the deep sediment AOA clades.

Many organisms possessing distinct electron transport, ion gradient generating and ATP synthase complexes that are differentially regulated under HHP [71, 86–88]. Interestingly, both NP-iota MAGs and two out of six NP-theta MAGs encode complete gene clusters for both the A-type ATPase found in neutrophilic AOA and V-type ATPase variant found in acidophilic/acidotolerant/piezotolerant archaea (and AOA) which is homologous to the proton/ion pumping ATPases from eukaryotes and enterococci (Fig. 4, 5) [89, 90]. The remaining NP-theta MAGs encode either the A-type or the V-type ATPase (although this could be attributed to the incompleteness of the MAGs). The V-type ATPase has been suggested to confer physiological advantages in high pressure environments by virtue of its proton-pumping function [89]. This would enable the maintenance of intracellular pH, which is disrupted by the accelerated release of protons from weak acids (such as carbonic acid) under HHP [91]. The presence of both ATPase variants is also observed in abysso/hadopelagic NP-gamma AOA lineages, while the deep marine NP-alpha encode only the V-type ATPase (Fig. 4,5 and Supplementary Information for further discussion) [89]. In contrast, all three NP-delta MAGs encode only the canonical A-type ATPase (Fig. 5), but intriguingly at least two of them seem to contain a partially duplicated NADH dehydrogenase (complex I) operon which could similarly be responsible to alleviate cytoplasm acidification (see Supplementary Information).

The cytoplasmic membrane is severely affected by HHP, which induces a tighter packing of the lipids and a transition to a gel state, resulting in a decrease in fluidity and permeability [92, 93]. The presence of a N-acetylneuraminic acid mutarotase (NanM) in NP-theta, NP-iota, NP-delta MAGs (Fig. 4, shared with a few abyssopelagic/hadal NP-gamma species) indicates the ability to acquire sialic acid [94]. This important component of glycoconjugates found on cell walls has multiple functions including concentrating water on cell surfaces [95] and regulating membrane permeability [96]. It can also enable the regulation of the thickness of the hydration layer surrounding the cell membrane [97], which could prevent system volume change and stabilize membrane protein complexes and membrane structure under pressure [97, 98], while also regulating membrane permeability [96]. Modification of the hydration layer properties has also been identified as a specific adaptation mechanism of the piezophilic archaeon *Thermococcus barophilus* [99].

An ABC-type branched-chain aminoacid transport system of the HAAT family (3.A.1.4) is present in all three sediment clades as well as in one NP-alpha MAG, sponge-associated and few other lineages of the NP-gamma clade, NS and non-AOA Thaumarchaea (Fig. 4, 5, Table S3). The uptake of amino acids has been interpreted earlier as indicative of the possibility of organic carbon utilization via enzymes participating in canonical amino acid biosynthesis pathways and present in all or most AOA (e.g. aspA, ilvA, ilvE, aspC glyA, GDH, ProDH) [10, 23, 37, 61]. Such mixotrophic strategies are also responsible for the enormous ecological success in oligotrophic environments of oceanic cyanobacterial lineages [100]. However, canonical amino acid degradation key enzymes such as amino acid hydroxylases, the branched-chain α-keto acid dehydrogenase complex or 2-ketoacid:ferredoxin oxidoreductases have not been detected in deep sea or sediment AOA clades, nor are their genomes particularly enriched in proteases (Fig. S4). On the other hand, a metabolic shift from expensive *de novo* biosynthesis of cellular materials (with proteins accounting for 56% of total energy investment in oxic environments) to recycling of exogenous or endogenous resources is observed in HHP-adapted microorganisms. [16, 18, 69, 85, 86, 101, 102]. Therefore, it seems more likely that amino acids are used for recycling, as suggested earlier for AOA by isotope tracer and NanoSIMS experiments with sediment and oceanic crust communities [16, 32, 33, 69], and as observed in the obligate piezophile *T. barophilus* and other facultative piezophiles [102–104]. Moreover, amino acids (mostly glutamate, proline and glutamine) can be accumulated as compatible solutes to ensure the stabilization of macromolecular structures upon pressure or temperature related stress [104, 105]. It cannot be ruled out though that amino acids are also used for replenishing the intracellular ammonia pool, with minimal production (if at all) of reducing equivalents (Fig. 5) (see Supplementary Information for detailed discussion).

The sediment clades and especially NP-theta encode an extended repertoire of enzymes for DNA and protein repair compared to other NP (details in Supplementary Information and Fig. 4, 5, S4). This is an indication of energy investment towards maintenance of cellular components, rectifying damage due to low turnover rates and cellular aging rather than active and fast growth [71, 82, 106]. This strategy together with dormancy is presumed to be responsible for persistence in subseafloor energy limited environments [107].

### Osmoregulation

All sediment clades do encode a putative bifunctional CTP:inositol-1-phosphate cytidylyltransferase/di-myo-inositol-1,3’-phosphate-1’-phosphate synthase (ipct/dipps), responsible for the synthesis of the compatible solute di-myo-inositol-1,3’-phosphate (DIP) [108]. The enzyme is also found in deep marine AOA clades [23] (Fig. 4, 5). Biosynthetic genes for this compatible solute have so far only been observed in organisms growing above 55^°^C, and have been extensively transferred between archaea and bacteria [109], making these AOA clades the first non-thermophilic organisms with the ability to synthesize this inositol derivative. Compatible solutes can confer resistance to various types of stress, so it is possible that this anionic solute has multiple roles in these polyextremophilic organisms [110] especially since no pathways for synthesis/uptake of known osmolytes such as mannosylglycerate, ectoine/hydroxyectoine or glycine/betaine were identified in the NP-theta, NP-iota, NP-delta MAGs (Fig. 4, 5, S3).

## Conclusions

Our comparative and phylogenomic analyses using 11 sediment-derived MAGs reported in this study, together with a large collection of AOA genomes with a broad phylogenetic and ecological distribution allowed us to study the evolution, diversification and adaptation mechanisms of AOA into deep marine environments. Based on phylogenomic analyses and different from earlier scenarios [111] we conclude that AOA from deep marine sediments evolved independently within (at least) three lineages. Although it seems that the ancestor of the *amo*A-NP-theta clade was pelagic and descendants of it occupied the deep marine sediments and the oxic subseafloor crust, it is likely that in the case of *amo*A-NP-iota and *amo*A-NP-delta, there was a transition from terrestrial ecosystems/freshwater sediment to marine sediments without having colonized the ocean water column first. Interestingly, all extended capacities and adaptations discussed in this manuscript are found to be combined in lineage *amo*A-NP-theta, which represents the most widely distributed and abundant clade ranging over different marine sediment layers, whereas the other two clades that share some of these features exhibit a more distinct distribution pattern.

All AOA adapted to marine sediments and investigated in this study are able to perform ammonia oxidation in combination with CO_2_ fixation like all other described AOA. In addition, all three lineages seem to be capable of utilising exogenous organic fermentation products that they convert into intermediates of their central carbon metabolism, a feature they share with pelagic AOA from the deep ocean and a few other AOA. This, together with the capability of taking up aminoacids, putatively for recycling into proteins or utilization of amine groups, would support growth in this extremely oligotrophic environment and contribute to organic nitrogen and carbon turnover in the sediments. In the absence of any components indicating increased capacity of amino acid degradation in these AOA we argue that recycling of amino acids rather than catabolism as otherwise suggested in [10, 23, 113] represents an advantageous and more plausible strategy for the sedimentary AOA clades. It is also a trait frequently observed in other sedimentary and crustal population groups to overcome the prohibitive energetic costs of *de novo* monomer biosynthesis [69]. Additionally, a broad repertoire of DNA and protein repair enzymes, seem to enable the deep sediment-adapted AOA to counteract the most severe consequences of cellular aging. An important feature shared with HHP-adapted deep marine clades is the presence of two ATPase complexes in *amo*A-NP-theta and *amo*A-NP-iota, with putatively opposing functions that would alleviate the effects of pH imbalance due to HHP, as well as the PMF-destabilizing effects of age-induced membrane leakage. These features shed light onto the mechanisms underlying AOA persistence in the benthic environments beneath the open ocean, from the surface sediments down to the underlying oceanic crust, and further consolidate the central role of these archaea in the global biogeochemical cycles.

## Author Contributions

CS, SLJ & TN conceived the study. RZ & SLJ sampled and processed the Atlantic sediments, while TN, HN, MH & YT sampled and processed the Pacific sediments. RZ, SSA & RP assembled the MAGs. RP performed the phylogenetic and comparative genomic analyses. MK annotated and analyzed the genomes. MK, RP, RZ, SSA, SLJ, CS & TN interpreted data. MK, RP & CS wrote the manuscript with contributions from TN, RZ, SLJ & SSA.

## Acknowledgements

We thank Philipp Weber for help with visualizations. The LABGeM (CEA/Genoscope & CNRS UMR8030), the France Génomique and French Bioinformatics Institute national infrastructures (funded as part of Investissement d’Avenir program managed by Agence Nationale pour la Recherche, contracts ANR-10-INBS-09 and ANR-11-INBS-0013) are acknowledged for support within the MicroScope annotation platform. We thank onboard scientists, officers and crews of RV *Yokosuka* and the manned submersible *Shinkai 6500* for their help and operation during the YK13-09 and YK13-12 cruises. We wish to acknowledge the entire scientific party and all crewmembers onboard *Joides Resolution* during IODP expedition 336, for their help and expertise. Especially, Co-chief scientists Katrina Edwards and Wolfgang Bach. We also wish to thank chief scientist of the CGB summer cruise 2014 Rolf Berger Pedersen for the sediment coring opportunity and thank Ingeborg Ørkland, Desiree Roerdink, Tamara Bumburger and Ingunn H Thorseth for their help with porewater extraction and nutrient analysis. This project was funded by FWF grant 27017 and ERC AdvGr TACKLE (695192). SLJ was funded by The Trond Mohn starting grant BFS2017REK03 and the Census of Deep Life phase V grant. TN was partially supported by JSPS KAKENHI Grant Number JP19H05684 within JP19H05679 (Post-Koch Ecology).

