## Supplementary Information, Methods and Figures for "Genomes of Thaumarchaeota from deep sea sediments reveal specific adaptations of three independently evolved lineages"

#### Site description of deep-sea seafloor samples

All sediment cores used in this study were retrieved from bathyal or abyssal depths. The two Pacific cores are fully oxic, whereas dissolved oxygen was not detectable in the middle part of NP_U1383E and the basal part of GS14-GC08 (Fig. S1). All except one (GC08-250 cm) of the metagenome sequencing datasets were generated from oxic sediment where nitrate was also detected in the porewater (Fig. 1b).

#### Amplicon sequencing and analysis

16S rRNA gene amplicons of the Atlantic sediments were prepared using primers Uni519f/806r and sequenced using an Ion Torrent Personal Genome Machine described previously [1]. The amplicons of the Pacific samples were prepared using a universal primer set of U530F and U907R and sequenced using Illumina MiSeq platform, following the procedure described in [2, 3]. To study the overall community structure of Thaumarchaeota, the sequencing data were processed as described elsewhere [1]. Briefly, the reads were quality-controlled and OTUs (97% nucleotide similarity threshold) were clustered using USEARCH and classified using CREST. For individual cores, OTUs classified as Nitrosopumilales were extracted from the OTU tables, and their clade affiliations were assigned based on their placement in the Nitrosopumilales 16S rRNA gene phylogenetic tree presented in [4].

#### High similarity between genomes from the Pacific and Atlantic sediments

Although the NP-iota MAGs (NPMR_S100_NP_iota_1 and YK1309_1N_S300_NP_iota) were assembled from metagenomic datasets of marine sediments in the Pacific and Atlantic oceans respectively, these MAGs showed 99 % ANI (Fig. S1), suggesting that these bins might represent different strains of the same species (prokaryotic species usually show > 95 % ANI among themselves [5, 6]). A similar pattern was observed between the Pacific bin YK1312_12N_S200_NP_theta and the Atlantic bin NPMR_S100_NP_theta_3, exhibiting > 97 % ANI.  It has been shown that prokaryotes with close phylogenetic affiliation inhabiting deep marine sediments and subsurface oceanic crust are commonly retrieved in distant geographic locations [7, 8], possibly due to the circulating seawater that allows the dispersion of subsurface and benthic microbial phyla  [9].

**Taxonomic placement of NPMR_NP_delta_1**

Based on the *amo*A tree MAG NPMR_NP_delta_1 clustered within the NP-theta clade (Fig. 2b) but the more robust phylogenomic analysis strongly suggests that it belongs to the NP-delta subclade (Fig 2a). We argue that the *amo*A gene in this MAG (in which *amo*AXC are the sole genes of a small contig) might represent a contamination considering that NP-delta and NP-theta have similar ecological distribution [4]; this study) and the phylogenomic tree reconstruction with 79 single-copy gene markers shows clearly that this MAG is placed within the NP-delta lineage.

**Notes on the evolution of AOA lineages from comparative genomics**

Interestingly, our estimation of 269 AOA-specific core families is close to the estimation of 289 protein families inferred to have been gained by the last common ancestor of AOA (Abby et al., 2020, submitted). We observed that extensive differential gene loss (and possibly gene acquisitions) have occurred at the origin of major AOA lineages as is attested by the great number of families shared between different combinations of protein families present in 2, 3 or 4 lineages only (Fig. 3, S2). Similarly, we found that each AOA lineage harbors between 500 to more than 3000 lineage-specific families suggesting a complex genomic evolution and a considerable number of gene acquisitions at the origin and during the diversification of AOA lineages, possibly associated to habitat adaptations [10–12] (Abby et al., 2020).

Based on genomic context analyses and considering the phylogenetic position of the NP-iota clade in the context of the scenario proposed by [13], it is plausible that the V-type ATPase was acquired by the common ancestor of NT/NP clades, followed by selective losses of one or the other in the resulting clades according to their environmental radiation. Genomic context analysis reveals that in NP-iota MAGs, the two operons are encoded next to each other, and flanked by the pyrI, pyrB and sulfT genes which are also the flanking genes of the V-type ATPase operon in NT and NP-alpha genomes/MAGs, as well as the A-type ATPase operon in all other NP clades [13]. In abysso/hadopelagic NP-gamma AOA, the V-type ATPase operon is located elsewhere in the genome (Table S3), implying an independent acquisition [13]. In any case, this distribution suggests that the acquisition of the proton-pumping ATPase variant was crucial for successful radiation into hadal high-pressure environments and raises intriguing possibilities about the ecophysiological potential of the ancestor of *Nitrosopumilales* (see evolution scenario above).

**Usage of exogenous organic compounds and high pressure adaptations**

The thaumarchaeal putative lactate racemase family enzyme has a 32% amino acid identity (1e-59) to the characterized LarA from *Lactobacillus plantarum*, and is a nickel-dependent enzyme activated by a maturation system [14] also found in the sediment AOA bins (Table S3). In lactobacilli, D-lactate is an important cell wall component conferring resistance to vancomycin [15], a glycopeptide antibiotic which inhibits cross-linking of N-acetylmuramic acid (NAM)/ N-acetylglucosamine (NAG) polymers. NAG/NAM is also a component of the thaumarchaeal cell surface, given the presence of NAG-utilizing enzymes in AOA (Table S3, [16]). The presence of Lar would enable the utilization of both lactate stereoisomers produced by the sediment fermentative community. The transport of lactate could be mediated by MIP family transporters (aquaporins) [17] common in AOA, as in lactobacilli. It has to be noted though that this enzyme belongs to a large superfamily of proteins with broad distribution in non-lactate utilizing organisms, probably catalyzing other racemization reactions [15, 18]. Phylogenetic analysis of the superfamily PF09861 of which LarA is a member reveals that the AOA homologs belong to a separate, but neighboring, cluster from the characterized LarA homologs from lactobacilli, leaving the question of the putative substrate open (Fig. S6)

The malate dehydrogenase (MDH) homologs encountered in AOA belong to the LDH-like MDH subgroup within the LDH/MDH superfamily of 2-ketoacid:NAD(P)-dependent dehydrogenases, as other archaeal MDHs [19]. In terms of primary sequence, quaternary structure and enzymatic properties, characterized archaeal homologs are between canonical MDHs and LDHs, possessing clear activity with oxaloacetate (as the former) but also able to utilize pyruvate (as the latter), while also exhibiting relaxed cofactor specificity (NADH or NADPH) [19, 20]. The active site architecture surrounding the universally conserved substrate binding residue (Arg171) resembles the environment found in canonical LDHs in AOA homologs, as in the characterized LDH-like MDH from *Ignicoccus islandicus* (Fig S5) [19]. In particular, while position 102 is occupied by an arginine and a neutral residue (methionine) is found in position 199, as in canonical MDHs, a threonine at position 246 and a histidine at position 68, typical LDH residues, may influence substrate selection and charge balance respectively (Fig. S5, residues highlighted in orange) [19, 20]. We therefore hypothesize that AOA homologs exhibit a broad substrate specificity and are in principle able to convert lactate to pyruvate, with the concomitant formation of NADH (cofactor preference for NADH is inferred by the presence of Asp54, highlighted in green in Fig. S4). However, whether this actually takes place *in vivo* necessitates enzymatic characterization.

No genes associated with the glycine cleavage system or choline/betaine degradation present in certain hadopelagic NP-gamma and NP-alpha lineages [21, 22] were identified in any of the sediment bins (Fig. S4).

Interestingly, a part of the NADH dehydrogenase (complex I) operon is duplicated in two out of three NP-delta MAGs (Fig. 5, Table S3) specifically genes nuoIJKML in NPMR_NP_delta_1 and nuoHIJKM in NPMR_NP_delta_3, bearing 85-95% amino acid identity. Unfortunately, these duplicated regions are in single contigs and therefore it is unclear whether the whole operon is duplicated. It is intriguing, however, that these regions contain the proton pumping subunits of complex I, raising the possibility that this could either be a mechanism to alleviate cytoplasm acidification under high pressure if the complex is running in the forward direction and pumping protons out, similar to the V-type ATPase. It should be noted here that Complex I is postulated to run in reverse in nitrifiers [23, 24]. Duplicated subunits or multiple copies of complex I are observed in various microorganisms including members of AOB and NOB, and are associated with increasing proton-pumping capacity or providing different electron flow options by operating in different directions, respectively [25–27]. Facultative piezophiles such as *Shewanella violacea* has been shown to encode distinct complexes of the respiratory chain (such as different versions of terminal oxidases) as an adaptation to growth at different pressure conditions [28].

**Usage of amino acids (AA)**

Enzymes participating in canonical amino acid biosynthesis pathways and present in almost all AOA (e.g. aspA, ilvA, ilvE, aspC glyA, GDH) could enable the utilization of imported amino acids such as Asp, Gly, Ser, Thr, Ile, Val, Leu, Phe, Tyr into their corresponding α-ketoacids or dicarboxylic acids by releasing a molecule of ammonia (Fig. 5, S4, Table S3). Only the catabolism of proline to glutamate by proline dehydrogenase (ProDH) and 1-pyrroline-5-carboxylate dehydrogenase (RocA), two enzymes with conserved gene synteny, is not widespread in AOA but present only in certain clades (Fig. 5, S4). Proline catabolism would generate reducing equivalents (Fig. 5) and result in the production of glutamate, which can be used to regulate the ammonia pool via the action of glutamate dehydrogenase (GDH) or would have a possible role in osmoregulation [29]. A type-III aminotransferase (Oat, CLUSTER_3296) encoded by the NP-theta, NP-iota MAGs could also participate in replenishing the intracellular ammonia stock by catalyzing the transamination between a variety of amino acids, mono- and diamines and α-ketoacids (Fig. 4, 5).

In obligate piezophiles such as *Thermococcus barophilus*, a drastic increase in AA requirements during HHP growth, even for those where biosynthesis pathways are present, was interpreted as a switch from energy-intensive AA biosynthesis towards recycling, in the context of the general metabolic response towards more efficient energy utilization under HHP [30–33]. A downregulation of AA biosynthesis pathways (especially glutamine, glutamate and costly aromatic AA), together with an upregulation of AA transport systems is observed during HHP growth in transcriptomic studies of facultative piezophiles *Desulfovibrio hydrothermalis* and *Desulfovibrio piezophilus*, While this was interpreted as solely resulting in glutamate accumulation in the cells (acting as a piezolyte), the accompanying shift in the energy metabolism of these organisms towards increased energy efficiency could also be taken as evidence of AA recycling. Moreover, in obligate piezophiles such as *Pyrococcus yayanosii*, certain AA biosynthesis pathways have been altogether lost [34], while *P. abyssi* also requires 9 amino acids for growth despite possessing biosynthesis pathways [35].

**Transporter complement of sediment clades**

All deep sediment clades encode the ion transporter repertoire typical for marine microorganisms (Table S3, Fig S4)([21–23, 36–39]: aquaporins (MIP family), small conductance mechanosensitive channels (MscS, accompanied by an absence of the large conductance MscL), NhaP-type $K^+(Na^+)/H^+$ antiporters (CPA1 family), while the low affinity but rapid uptake Trk family transporter is only encoded by the NP-theta and NP-gamma clades. Additionally, NP-theta, NP-iota, hadopelagic NP-gamma and deep marine NP-alpha lineages encode ArsB family $Na^+/H^+$ antiporters.

**Motility and attachment**

Only the NP-delta bins encode a complete repertoire for archaellum assembly and chemotaxis, while both NP-delta and NP-theta clades encode putative Type IV pili assembly clusters (Fig 4, 5, S4, Table S3). The absence of an archaellum in the clades most adapted to this habitat is not surprising, as this is one of the most sensitive apparatuses and processes to high hydrostatic pressure, while also extremely energy demanding and therefore rarely occurring in sediment lineages [31, 40, 41]. Conversely, the capacity for attachment indicated by the Type IV pili could provide the possibility to adhere to energy-rich particles, and has been implicated in starvation survival strategies [31]. It would appear therefore that active migration (at a huge energy cost) in the sediment column would be an option only for NP-delta, while NP-theta representatives are well adapted to a sedentary lifestyle.

**Information processing systems, DNA and protein repair**

DNA depurination is the most common source of age-induced DNA damage in marine sediments [42, 43]. The main DNA repair systems in the three sediment lineages seem to be double strand break repair (DBSB) via homologous recombination (HR), and base excision repair (BER), while key NER enzymes such as XPD and XPB/Bax1 are missing, as in most *Ca*. Nitrosopumilales & *Ca.* Nitrosotaleales (Table S3, Fig 4, 5 & S4). While all essential proteins for HR are present in NP-theta and NP-iota, NP-delta bins lack a homolog of the Hef helicase domain protein involved in repair of stalled replication forks [44]. However, functional redundancy between Hef and Hjc (Holliday junction resolvase), present in all AOA, has been observed in *H. volcanii*, ensuring that all sediment bins are able to process Holliday junctions and restart collapsed replication forks [45].

The arsenal of monofunctional and bifunctional DNA glycosylases and endonucleases involved in base excision repair (BER) [46], the pathway responsible for the repair of modified (oxidized, alkylated or deaminated) or mismatched bases is present, with some differences, in all sediment clades (Fig. 4, 5, S4). For example, while NP-theta bins encode uracil DNA glycosylases of families 4&5 (Udg), a methylpurine/alkyladenine-DNA glycosylase (Mpg), a 3-methyladenine DNA glycosylase (AlkA) and an 8-oxoguanine (8-oxoG) DNA glycosylase (Ogg1), NP-delta bins encode only family 5 Udg and AlkA glycosylases, while NP-iota bins encode family 4 Udg, Mpg and Ogg1 glycosylases as in hadal NP-α lineages [22]. Additionally, they all encode, as all AOA, an EndoIII/Nth homolog, which has both 8-oxoG DNA glycosylase/AP (apurinic or apyrimidinic site) lyase activities. It is plausible that NP-theta bins encoding four different DNA glycosylases are better equipped at recognizing various types of DNA damage than other NP clades. However, it should be noted that while all superfamilies of glycosylases exhibit substrate specificity, e.g. Udg for uracil pairs [47], Ogg1 for 8-oxoG, AlkA and Mpg for methylated and alkylated bases [48], there is a substrate overlap among superfamily members (e.g. among AlkA and Mpg), so in principle all clades have the capability to recognize the basic types of damaged bases.

The UvrABC system of NER [49, 50], present in AOA, is present only in NP-delta and one NP-theta bin and is also absent from the hadalopelagic and deep marine NP representatives (Fig. 4, 5, Table S3). Interestingly, the Uvr system has been implicated in transcription-coupled repair in halophilic archaea [51].

While all clades encode the single subunit family B polymerase PolB1, no family D polymerase subunits (DP1, DP2) were detected in NP-iota (Fig. 5, S4). The absence of PolD is also observed in hadal lineages belonging to the NP-alpha clade (erroneously referred to as PolB by [22]) as well as Ca. *Nitrosocaldales* [10, 52]. This provides even stronger support to the hypothesis that PolB1 is the replicative polymerase in Thaumarchaea, performing both leading and lagging strand synthesis as in *S. solfataricus* [53]. This would indicate that PolD in those Thaumarchaea where present has a secondary role in repair, in reversal to the situation observed in Euryarchaeota where PolD is the replicative polymerase while the recognition and complete inhibition by deaminated bases of PolB (PolB3 group) implicated it in repair pathways [54–56]. In the absence of any functional data for the thaumarchaeal polymerases, it is unclear what role PolD plays in repair and what kind of impediment is caused by its absence. Data from euryarchaeal homologs however indicate that this family also stalls upon encountering deaminated bases and can inhibit BER, and could therefore participate in replication-associated repair [54].

NP-theta and NP-iota bins encode a protein-L-isoaspartate carboxylmethyltransferase homolog (pcm), responsible for the repair of D-aspartyl residues in proteins, also found in NP-gamma and NP-eta clades (Fig. 4, 5)[36]. Spontaneous aminoacid racemization is one the most important causes of protein damage in the subsurface marine environments due to the low turnover rates [57], indicating an energy investment towards cell maintenance.

Adaptation to the low temperatures in the deep sediments is assisted by homologs of the cold-shock protein CspC [58], present in the NP-theta and NP-gamma clades, and the cold-shock DEAD-box protein A (CshA), present in all NP (Fig. 4, 5) [59].

### Supplementary Methods Information

#### Porewater geochemistry of Pacific sediment samples

Sample processing of Pacific sediment cores: Upon recovery on board, sediment cores were kept in a cold room (4ºC) prior to sample processing. Overlying water was gently sampled and filtrated for geochemical analyses, and then sediment cores were sliced horizontally into 1 to 5cm thickness of sediments. For porewater extraction, sediment samples were immediately centrifuged at 2600 g for 5 min. Extracted porewater were filtered with a 0.45-µm membrane filter and stored at -20° C. Samples for molecular analyses were stored at -80ºC. Porewater geochemistry of the Pacific sediment samples is described in Supplementary Information.

Dissolved oxygen (DO) concentrations in the sediment cores were measured onboard immediately after core recovery using a planar optode oxygen sensor Fibox 3 (PreSens, Regensburg, Germany). The sensor spots were attached to the inside of the transparent polycarbonate core liner tube and oxygen concentrations were measured from the outside [60]. The sensors were calibrated with air-saturated and oxygen-free seawater before measurements.

Nutrient concentrations were measured with a continuous-flow analyser (BL-Tech QUAATRO 2-HR system) [61] onboard. The precision of the phosphate, nitrate, nitrite, and ammonium measurements, based on duplicate measurements, was ± 0.17%, ± 0.17%, ± 0.16%, and ± 0.38%, respectively. When a nutrient concentration exceeded the calibration range, the filtrated porewater was diluted with nutrient-free seawater and were measured again to reduce the concentration to within the calibration range.

**DNA extraction, library construction and sequencing**

DNA for metagenomic sequencing from the Atlantic sediment samples was extracted from ~7 g sediment (~0.7 g sediment in 10 individual lysis tubes) using PowerLyze Soil DNA Isolation Kit (MoBio Laboratories) following the manufacturer's instructions, except for the following minor modification: the lysing tubes were incubated in water bath of 60°C for 15 min prior to beading beating at the highest speed (grade of 6) for 45 seconds on the MP machine. The DNA extracts were iteratively eluted from the 10 spin columns into 100 µL of $ddH_2O$ for further analysis.

DNA was sheared into 400 bp fragments using Covaris, and libraries were constructed using a Nextera DNA Flex Library Prep kit (Illumina). Metagenomic libraries were sequenced (2×150 bp paired-end) by an Illumina Hiseq 2500 sequencer at the Vienna Biocenter Core Facilities GmbH (Vienna, Austria).

DNA extraction, purification and shotgun metagenomic library construction from the Pacific sediment samples were conducted as described previously [3]. Briefly, from approximately 5 g of the frozen sediment DNA was extracted using DNeasy PowerMax Soil Kit (QIAGEN) and further purified with DNA Clean up MagExtractorTM –PCR & Gel Clean up- (TOYOBO). Then, shotgun metagenomic libraries were constructed using KAPA Hyper Prep Kit from 1 ng DNA or less. The metagenomic sequence libraries were analyzed using Illumina HiSeq2500 with rapid mode (250 bp paired-end).

**Assembly and genome binning of Atlantic sediments**

The sequencing data were processed with Trimmomatic v.0.36 [62] to remove illumina adapters and low quality reads ("SLIDINGWINDOW:10:25"). The quality-controlled reads from the eight samples were de novo co-assembled into contigs using Megahit v.1.1.2 [63] with the k-mer length varying from 27 to 117. Contigs larger than 1000 bp were into automatically binned with MaxBin2 v2.2.5 [64] using the default parameters. The quality of the obtained genome bins was assessed using CheckM v.1.0.7 [65] with the option "lineage_wf", which uses lineage-specific sets of single-copy genes to estimate completeness and contamination and assigns contamination to strain heterogeneity if amino acid identity is >90%. Genome bins of >50% completeness were manually refined using the gbtools [66] based on the GC content, taxonomic assignments, and differential coverages in different samples. Coverages of contigs in each sample were determined by mapping trimmed reads onto the contigs using BBMap v.37.61 [67]. Taxonomy of contigs were assigned according to the taxonomy of the single-copy marker genes in contigs identified using a script modified from blobology [68] and classified by BLASTn. SSU rRNA sequences in contigs were identified using Barrnap (Seeman 2015, Github), and classified using VSEARCH with the SILVA 132 release [69] as the reference.

To improve the quality of the Thaumarchaeota genomes, we recruited reads from highest-abundance-sample (i.e. highest genome coverage) using BBMap as described above, and the recruited reads were re-assembled using SPAdes v.3.12.0 [70]. After removal of contigs shorter than 1 kb, the resulting scaffolds were visualized and re-binned muanlly using gbtools [66] as described above. The quality of the resulting Thaumarchaeota genomes were checked using the CheckM v.1.0.7 "lineage_wf" command again, based on the Thaumarchaoeta marker gene set (automatically selected by CheckM).

**Metagenomic assembly and binning of Pacific samples**

The reads of the 6 metagenomic samples from deep marine sediments of the Pacific Ocean were quality trimmed and illumina adapters were removed using Trimmomatic [62]. Low-complexity homopolymeric reads (sequences with more than 80% of a single nucleotide) were removed with PRINSEQ [71]. The quality trimmed reads of each metagenomic sample were assembled independently using MEGAHIT [63]. The trimmed reads of all Pacific metagenomes were then mapped back onto the 6 different assemblies using the bowtie2 tool [72].

The contigs of the six metagenomes were binned with CONCOCT [73], MetaBAT [74] and MaxBin 2 [64] followed by contig dereplication and binning optimization using the DAS tool [75]. Completeness and contamination of bins were evaluated through single copy-marker gene comparison with CheckM (same parameters as above) [65]. Prodigal [76] was employed with default parameters to predict proteins of optimized bins.

**Phylogeny of *amo*A sequences**

The nucleotide *amo*A sequences of the MAGs reported in this study were retrieved via BLASTN searches (E-value $10^{-10}$) using the *amo*A sequence of *Nitrosopumilus ureiphilus* (KX950756.1). The *amo*A sequences were incorporated into the curated alignment of *amo*A genes reported by [77], using MAFFT v7 ("--add" parameter) [78] followed by the manual inspection of the alignment.

The maximum likelihood phylogenetic tree of *amo*A sequences was reconstructed using IQTREE (v2.0-rc1) [79] with a GTR+F+I+G model of sequence evolution using a constrained tree search ("-g" parameter) based on the phylogenetic tree reported in [77], and 1,000 ultrafast bootstrap replicates.

**Annotation and comparative genomics analysis**

In addition to the 11 AOA MAGs reported in this work (Table 1), we downloaded 18 completely sequenced AOA genomes, 13 nearly complete genomes and 43 metagenome-assembled or single-amplified genomes from NCBI or IMG database (date: December 2019). In total, our genome collection was composed of 85 genomes (77 AOA plus 8 non-AOA Thaumarchaeota). The complete list of genomes is provided in Table S1. All MAGs and SAGs collected from public databases are more than 70 % complete and have less than 5% contamination, except for Marine Group I thaumarchaeote SCGC RSA3 and Marine Group I thaumarchaeote SCGC AB-629-I23 which are more than 90 % complete and have less than 10 % contamination. Prodigal [76] was used with default parameters to predict protein sequences when this information was not provided. Annotation of the MAGs assembled in this study was performed automatically using the Microscope annotation platform from Genoscope [80], followed by extensive manual curation.

**ANI comparisons**

Pairwise average nucleotide identity comparisons between MAGs were performed using the ANI script from the enveomics collection [81].

**Sequence alignment**

Primary sequences were aligned with Mafft [82] and visualized with BOXSHADE (ExPASy).

**Data availability**

Raw reads from metagenomic sequencing as well as 16S and *amo*A amplicons have been submitted to NCBI under project accession numbers PRJNA489438, PRJNA529480 (Atlantic samples) and PRJDB9793 (Pacific samples). Assemblies of MAGs reported in this study are available at NCBI under accession numbers (pending) and available in the Microbial Genome Annotation & Analysis Platform Microscope (https://mage.genoscope.cns.fr/microscope/home/index.php) [80].

**Supplementary Figures**

**Supplementary figure 1 (S1).** Porewater profiles of oxygen and nitrate in the sediment cores used in this study. In the two Pacific cores (YK1309-1N and YK1312-12N), the dashed lines denote the sediment-water interface. Note in the two Atlantic cores the sediment-water interfaces were not properly recovered by the piston/gravity coring. Different axis scales were used for different cores. Data of NP-U1383E were from [1], and GS14-GC08 from [83].

**Supplementary figure 2 (S2).** Pairwise average nucleotide identity calculations among the the deep marine sediments-derived MAGs. The archaeon CSP1 MAG was included in the analyses to have all the representatives of the NP-delta clade.

**Supplementary figure 3 (S3).** ML Phylogenetic tree of 2HADH sequences.
A dehydrogenase protein cluster comprising 14 sequences (colored in blue in the phylogenetic tree) was added using MAFFT v7 [82] ("mafft-linsi –add") to the structure-based reference alignment of 2HADH utilized by [84]. The alignment was trimmed in trimmAl ("trimal -gt 0.2") and a ML phylogenetic tree was calculated using IQTREE (v2.0-rc1) [79] with a GTR+F+I+G model and 1,000 ultrafast bootstrap replicates. Yellow circles represent nodes with 100 % bootstrap support. The scale bar indicates the number of amino acid substitutions per site.

**Supplementary figure 4 (S4).** Extended heatmap depicting the distribution and abundance of genes involved in the main functional categories discussed in the text. Abbreviations: nit2, nitrilase/omega-amidase; ureA, urease subunit gamma; mco1, multicopper oxidase family 1; fdh, formate dehydrogenase; larA, lactate racemase; pgi, phosphoglucose isomerase; proDH, proline dehydrogenase; rocA, 1-pyrroline-5-carboxylate dehydrogenase; oat, putative ornithine--oxo-glutarate aminotransferase/class III aminotransferase; ggt, g-glutamyl transpeptidase; kal, 3-aminobutyryl-CoA ammonia lyase; kat, putative 3-aminobutyryl-CoA aminotransferase; argD, acetylornithine transaminase; serC, serine-puryvate aminotransferase; ilvA, threonine/serine ammonia-lyase; ilvE, branched-chain-amino-acid transaminase; aspC, aspartate/tyrosine/aromatic aminotransferase; gvtTPH, glycine cleavage system proteins T/P/H; metH, methionine synthase II (cobalamin-independent); metE, methionine synthase I (cobalamin-dependent); tbp, TATA-box binding protein; rpoS54, AAA family ATPase/RNA polymerase sigma factor 54 interaction domain; phr, photolyase; polD2, DNA polymerase D, large subunit DP2; uvrABC, the Uvr excision repair system endonucleases ABC; hef, Hef/FANCM/Mph1-like helicase; udg4/5, Uracil DNA glycosylase family 4/5; mpg, methylpurine/alkyladenine-DNA glycosylase; ogg1, 8-oxoguanine DNA glycosylase; alkA, DNA-3-methyladenine glycosylase; tag, 3-methyladenine DNA glycosylase; endoV, endonuclease V; POP4,

RNase P/RNase MRP subunit p29, uspA, universal stress protein A; pcm, protein-L-isoaspartate carboxylmethyltransferase; ipct/dipps, bifunctional CTP:inositol-1-phosphate cytidylyltransferase/di-myo-inositol-1,3′-phosphate-1′-phosphate synthase; cspC, cold-shock protein A; cshA, cold-shock DEAD-box protein A; LLM, luciferase-like monooxygenase family protein; nanM, N-acetylneuraminic acid mutarotase; flaK, archaeal preflagellin peptidase FlaK; cheY, chemotaxis response regulator CheY; cheAB, chemotactic sensor histidine kinase cheA & methylesterase cheB; XerD/XerC family integrases; Protease and transporter classes can be found in Table S4. All locus tags and cluster information are in Supplementary tables 3 & 4.

**Supplementary figure 5 (S5).** Sequence alignment of lactate dehydrogenase (LDH) and malate dehydrogenase (MDH) homologs, shaded in yellow and green, respectively. Conserved residues are shaded in black and grey. The universally conserved substrate binding Arg171 in the LDH/MDH superfamily is indicated in red. Residues important for substrate discrimination and active site architecture are shaded in orange and discussed in the text. The residue determining the cofactor specificity is shaded in green. Residue numbering refers to the LDH numbering as in [19].

Primary sequences from the following organisms were used to generate the alignment (Uniprot accession numbers in parentheses): Gster, *Geobacillus stearothermophilus* (P00344); Tth, *Thermus thermophilus* (Q5SJA1); Blon, *Bifidobacterium longum* (E8ME30); Ctep, *Chlorobaculum tepidum* (P80039); Nvie, *Nitrososphaera viennensis* (A0A060HG74); Nmar, *Nitrosopumilus maritimus* (A9A450); Nkor, *Nitrosarchaeum koreense* (F9CUM5); Nbrev, *Ca.* Nitrosopelagicus brevis (A0A0A7V4F4); Nuzo, *Ca.* Nitrosotenuis uzonensis (V6AR53); Mjan, *Methanocaldococcus jannaschii* (Q60176); Iisl, *Ignicoccus islandicus* (A0A0U3FQH7); Msed, *Metallosphaera sedula* (A4YDY0). Sequences from the marine sediment AOA MAGs reported in this study are in bold and their locus tags can be found in Table S3.

**Supplementary figure 6 (S6).** Phylogenetic tree of lactate racemase sequences. A total of 1813 proteins of the PF09861 superfamily, 19 LarA sequences reported by [15] (colored in blue) and 7 putative LarA sequences identified in the MAGs reported in this study (colored in red) were aligned using MAFFT v7 (FFT-NS-2 strategy)[85]. The alignment output was used to construct an approximately-maximum-likelihood phylogenetic tree using FastTree v2.1.11 with default parameters [86].

**Description of Supplementary Tables**

**Table S1**: List and description of genomes (complete, MAGs and SAGs) used in the study

**Table S2**: List of archaeal single copy marker genes from Rinke et al. 2013 used in the concatenated alignment for the generation of the phylogenomic tree in Fig. 2.

**Table S3**: Description of protein families generated from the dataset.

**Table S4**: Locus tags and associated annotation of central metabolic pathways of the MAGs reported in this study.

30. Vannier P, Michoud G, Oger P, Marteinsson VP, Jebbar M. Genome expression of Thermococcus barophilus and Thermococcus kodakarensis in response to different hydrostatic

pressure conditions. *Res Microbiol* 2015; **166**: 717–725.

31. Lever MA, Rogers KL, Lloyd KG, Overmann J, Schink B, Thauer RK, et al. Life under extreme energy limitation: a synthesis of laboratory- and field-based investigations. *FEMS Microbiol Rev* 2015; **39**: 688–728.
32. Amrani A, Bergon A, Holota H, Tamburini C, Garel M, Ollivier B, et al. Transcriptomics Reveal Several Gene Expression Patterns in the Piezophile Desulfovibrio hydrothermalis in Response to Hydrostatic Pressure. *PLoS One* 2014; **9**: e106831.
33. Amrani A, van Helden J, Bergon A, Aouane A, Ben Hania W, Tamburini C, et al. Deciphering the adaptation strategies of Desulfovibrio piezophilus to hydrostatic pressure through metabolic and transcriptional analyses. *Environ Microbiol Rep* 2016; **8**: 520–526.
34. Michoud G, Jebbar M. High hydrostatic pressure adaptive strategies in an obligate piezophile Pyrococcus yayanosii. *Sci Rep* 2016; **6**: 27289.
35. Watrin L, Martin-Jezequel V, Prieur D. Minimal Amino Acid Requirements of the Hyperthermophilic Archaeon Pyrococcus abyssi, Isolated from Deep-Sea Hydrothermal Vents. *Appl Environ Microbiol* 1995; **61**: 2069–2069.
36. Ngugi DK, Blom J, Alam I, Rashid M, Ba-Alawi W, Zhang G, et al. Comparative genomics reveals adaptations of a halotolerant thaumarchaeon in the interfaces of brine pools in the Red Sea. *ISME J* 2015; **9**: 396–411.
37. Offre P, Kerou M, Spang A, Schleper C. Variability of the transporter gene complement in ammonia-oxidizing archaea. *Trends Microbiol* 2014; **22**: 665–675.
38. Santoro AE, Dupont CL, Richter RA, Craig MT, Carini P, McIlvin MR, et al. Genomic and proteomic characterization of 'Candidatus Nitrosopelagicus brevis': An ammonia-oxidizing archaeon from the open ocean. *Proc Natl Acad Sci U S A* 2015; **112**: 1173–8.
39. Bayer B, Vojvoda J, Offre P, Alves RJ, Elisabeth NH, Garcia JA, et al. Physiological and genomic characterization of two novel marine thaumarchaeal strains indicates niche differentiation. *ISME J* 2016; **10**: 1051–1063.
40. Bartlett DH. Pressure effects on in vivo microbial processes. *Biochim Biophys Acta - Protein Struct Mol Enzymol* 2002; **1595**: 367–381.
41. Tully BJ, Heidelberg JF. Potential Mechanisms for Microbial Energy Acquisition in Oxic Deep Sea Sediments. *Appl Environ Microbiol* 2016; AEM.01023-16.
42. Hoehler TM, Jørgensen BB. Microbial life under extreme energy limitation. *Nat Rev Microbiol* 2013; **11**: 83–94.
43. Jørgensen BB, Marshall IPG. Slow Microbial Life in the Seabed. *Ann Rev Mar Sci* 2016; **8**: 311–332.
44. Ishino Y, Narumi I. DNA repair in hyperthermophilic and hyperradioresistant microorganisms. *Curr Opin Microbiol* 2015; **25**: 103–112.
45. White MF, Allers T. DNA repair in the archaea—an emerging picture. *FEMS Microbiol Rev* 2018.
46. Grasso S, Tell G. Base excision repair in Archaea: back to the future in DNA repair. *DNA Repair* 2014; **21**: 148–157.
47. Lucas-Lledo JI, Maddamsetti R, Lynch M. Phylogenomic Analysis of the Uracil-DNA Glycosylase Superfamily. *Mol Biol Evol* 2011; **28**: 1307–1317.
48. Denver DR. An Evolutionary Analysis of the Helix-Hairpin-Helix Superfamily of DNA Repair Glycosylases. *Mol Biol Evol* 2003; **20**: 1603–1611.
49. Morita R, Nakane S, Shimada A, Inoue M, Iino H, Wakamatsu T, et al. Molecular Mechanisms of the Whole DNA Repair System: A Comparison of Bacterial and Eukaryotic Systems. *J Nucleic Acids* 2010; **2010**: 1–32.
50. Truglio JJ, Croteau DL, Van Houten B, Kisker C. Prokaryotic Nucleotide Excision Repair:

The UvrABC System. *Chem Rev* 2006; **106**: 233–252.

51. Stantial N, Dumpe J, Pietrosimone K, Baltazar F, Crowley DJ. Transcription-coupled repair of UV damage in the halophilic archaea. *DNA Repair (Amst)* 2016; **41**: 63–68.
52. Daebeler A, Herbold CW, Vierheilig J, Sedlacek CJ, Pjevac P, Albertsen M, et al. Cultivation and Genomic Analysis of "Candidatus Nitrosocaldus islandicus," an Obligately Thermophilic, Ammonia-Oxidizing Thaumarchaeon from a Hot Spring Biofilm in Graendalur Valley, Iceland. *Front Microbiol* 2018; **9**.
53. Yan J, Beattie TR, Rojas AL, Schermerhorn K, Gristwood T, Trinidad JC, et al. Identification and characterization of a heterotrimeric archaeal DNA polymerase holoenzyme. *Nat Commun* 2017; **8**: 15075.
54. Abellón-Ruiz J, Ishino S, Ishino Y, Connolly BA. Archaeal DNA Polymerase-B as a DNA Template Guardian: Links between Polymerases and Base/Alternative Excision Repair Enzymes in Handling the Deaminated Bases Uracil and Hypoxanthine. *Archaea* 2016; **2016**: 1–8.
55. Kushida T, Narumi I, Ishino S, Ishino Y, Fujiwara S, Imanaka T, et al. Pol B, a Family B DNA Polymerase, in *Thermococcus kodakarensis* is Important for DNA Repair, but not DNA Replication. *Microbes Environ* 2019; **34**: 316–326.
56. Makarova KS, Krupovic M, Koonin E V. Evolution of replicative DNA polymerases in archaea and their contributions to the eukaryotic replication machinery. *Front Microbiol* 2014; **5**: 354.
57. Mhatre SS, Kaufmann S, Marshall IPG, Obrochta S, Andrèn T, Jørgensen BB, et al. Microbial biomass turnover times and clues to cellular protein repair in energy-limited deep Baltic Sea sediments. *FEMS Microbiol Ecol* 2019.
58. Lopez-Garcia P, Zivanovic Y, Deschamps P, Moreira D. Bacterial gene import and mesophilic adaptation in archaea. *Nat Rev Microbiol* 2015; **13**: 447–456.
59. Français M, Carlin F, Broussolle V, Nguyen-Thé C. Bacillus cereus cshA Is Expressed during the Lag Phase of Growth and Serves as a Potential Marker of Early Adaptation to Low Temperature and pH. *Appl Environ Microbiol* 2019; **85**.
60. Hiraoka S, Hirai M, Matsui Y, Makabe A, Minegishi H, Tsuda M, et al. Microbial community and geochemical analyses of trans-trench sediments for understanding the roles of hadal environments. *ISME J* 2020; **14**: 740–756.
61. Nomaki H, Arai K, Suga H, Toyofuku T, Wakita M, Nunoura T, et al. Sedimentary organic matter contents and porewater chemistry at upper bathyal depths influenced by the 2011 off the Pacific coast of Tohoku Earthquake and tsunami. *J Oceanogr* 2016; **72**: 99–111.
62. Bolger AM, Lohse M, Usadel B. Trimmomatic: a flexible trimmer for Illumina sequence data. *Bioinformatics* 2014; **30**: 2114–2120.
63. Li D, Liu C-M, Luo R, Sadakane K, Lam T-W. MEGAHIT: an ultra-fast single-node solution for large and complex metagenomics assembly via succinct de Bruijn graph. *Bioinformatics* 2015; **31**: 1674–1676.
64. Wu Y-W, Simmons BA, Singer SW. MaxBin 2.0: an automated binning algorithm to recover genomes from multiple metagenomic datasets. *Bioinformatics* 2016; **32**: 605–607.
65. Parks DH, Imelfort M, Skennerton CT, Hugenholtz P, Tyson GW. CheckM: assessing the quality of microbial genomes recovered from isolates, single cells, and metagenomes. *Genome Res* 2015; **25**: 1043–1055.
66. Seah BKB, Gruber-Vodicka HR. gbtools: Interactive Visualization of Metagenome Bins in R. *Front Microbiol* 2015; **6**.
67. Bushnell B. BBMap: a fast, accurate, splice-aware aligner. *Joint Genome Institute, department of energy* . 2014.

68. Kumar S, Jones M, Koutsovoulos G, Clarke M, Blaxter M. Blobology: exploring raw genome data for contaminants, symbionts and parasites using taxon-annotated GC-coverage plots. *Front Genet* 2013; **4**.
69. Quast C, Pruesse E, Yilmaz P, Gerken J, Schweer T, Yarza P, et al. The SILVA ribosomal RNA gene database project: improved data processing and web-based tools. *Nucleic Acids Res* 2012; **41**: D590–D596.
70. Bankevich A, Nurk S, Antipov D, Gurevich AA, Dvorkin M, Kulikov AS, et al. SPAdes: A New Genome Assembly Algorithm and Its Applications to Single-Cell Sequencing. *J Comput Biol* 2012; **19**: 455–477.
71. Schmieder R, Edwards R. Quality control and preprocessing of metagenomic datasets. *Bioinformatics* 2011; **27**: 863–864.
72. Langmead B, Salzberg SL. Fast gapped-read alignment with Bowtie 2. *Nat Methods* 2012.
73. Alneberg J, Bjarnason BS, de Bruijn I, Schirmer M, Quick J, Ijaz UZ, et al. Binning metagenomic contigs by coverage and composition. *Nat Methods* 2014; **11**: 1144–1146.
74. Kang DD, Froula J, Egan R, Wang Z. MetaBAT, an efficient tool for accurately reconstructing single genomes from complex microbial communities. *PeerJ* 2015; **3**: e1165.
75. Sieber CMK, Probst AJ, Sharrar A, Thomas BC, Hess M, Tringe SG, et al. Recovery of genomes from metagenomes via a dereplication, aggregation and scoring strategy. *Nat Microbiol* 2018; **3**: 836–843.
76. Hyatt D, Chen GL, LoCascio PF, Land ML, Larimer FW, Hauser LJ. Prodigal: Prokaryotic gene recognition and translation initiation site identification. *BMC Bioinformatics* 2010.
77. Alves RJE, Minh BQ, Urich T, von Haeseler A, Schleper C. Unifying the global phylogeny and environmental distribution of ammonia-oxidising archaea based on amoA genes. *Nat Commun* 2018; **9**: 1517.
78. Katoh K, Standley DM. MAFFT multiple sequence alignment software version 7: improvements in performance and usability. *Mol Biol Evol* 2013; **30**: 772–780.
79. Minh BQ, Schmidt HA, Chernomor O, Schrempf D, Woodhams MD, von Haeseler A, et al. IQ-TREE 2: New Models and Efficient Methods for Phylogenetic Inference in the Genomic Era. *Mol Biol Evol* 2020; **37**: 1530–1534.
80. Vallenet D, Calteau A, Cruveiller S, Gachet M, Lajus A, Josso A, et al. MicroScope in 2017: an expanding and evolving integrated resource for community expertise of microbial genomes. *Nucleic Acids Res* 2017; **45**: D517–D528.
81. Rodriguez-R L, Konstantinidis K. The enveomics collection: a toolbox for specialized analyses of microbial genomes and metagenomes. 2016.
82. Kazutaka Katoh DMS. MAFFT Multiple Sequence Alignment Software Version 7: Improvements in Performance and Usability. *Mol Biol Evol* 2013; **30**: 772.
83. Zhao R, Mogollón JM, Abby SS, Schleper C, Biddle JF, Roerdink D, et al. In situ growth of anammox bacteria in subseafloor sediments. *bioRxiv* 2019; 729350.
84. Matelska D, Shabalin IG, Jabłońska J, Domagalski MJ, Kutner J, Ginalski K, et al. Classification, substrate specificity and structural features of D-2-hydroxyacid dehydrogenases: 2HADH knowledgebase. *BMC Evol Biol* 2018; **18**: 199.
85. Katoh K, Standley DM. MAFFT: iterative refinement and additional methods. *Methods Mol Biol* 2014; **1079**: 131–146.
86. Price MN, Dehal PS, Arkin AP. Fasttree: Computing large minimum evolution trees with profiles instead of a distance matrix. *Mol Biol Evol* 2009.

Figure S1

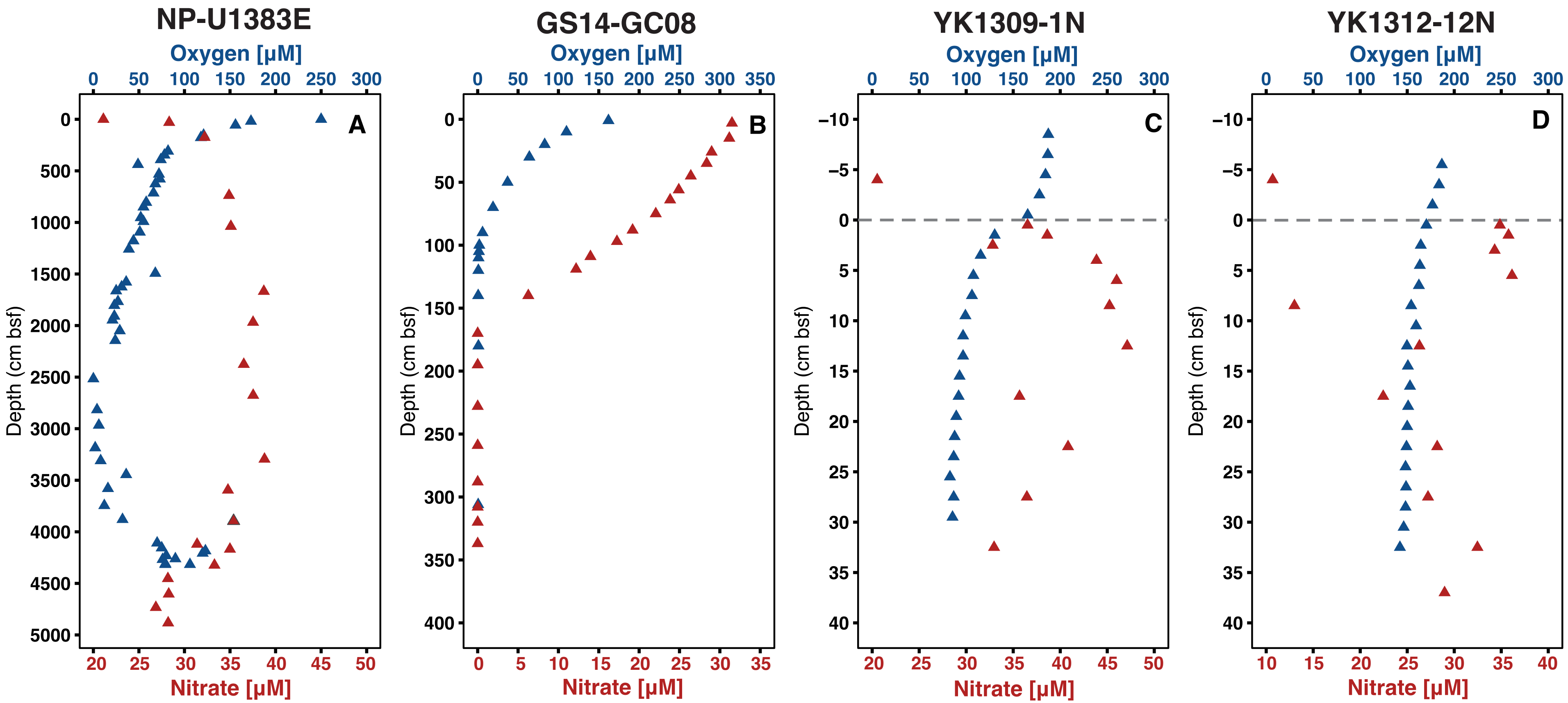
NP-U1383E
GS14-GC08
YK1309-1N
YK1312-12N
Oxygen [µM]
A
B
C
D
Depth (cm bsf)
Nitrate [µM]

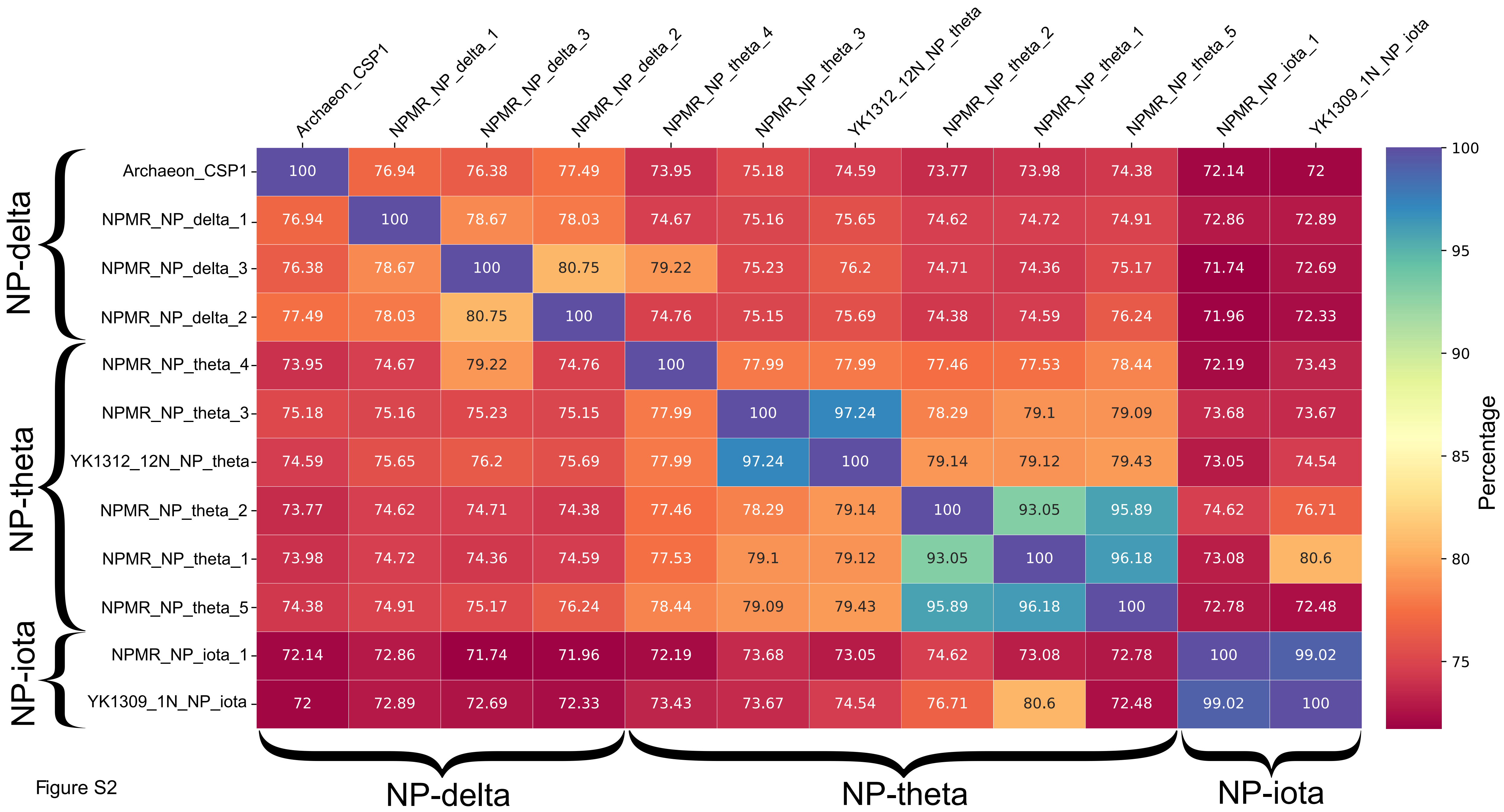
Archaeon_CSP1
NPMR_NP_delta_1
NPMR_NP_delta_3
NPMR_NP_delta_2
NPMR_NP_theta_4
NPMR_NP_theta_3
YK1312_12N_NP_theta
NPMR_NP_theta_2
NPMR_NP_theta_1
NPMR_NP_theta_5
NPMR_NP_iota_1
YK1309_1N_NP_iota
Archaeon_CSP1 100 76.94 76.38 77.49 73.95 75.18 74.59 73.77 73.98 74.38 72.14 72
NPMR_NP_delta_1 76.94 100 78.67 78.03 74.67 75.16 75.65 74.62 74.72 74.91 72.86 72.89
NPMR_NP_delta_3 76.38 78.67 100 80.75 79.22 75.23 76.2 74.71 74.36 75.17 71.74 72.69
NPMR_NP_delta_2 77.49 78.03 80.75 100 74.76 75.15 75.69 74.38 74.59 76.24 71.96 72.33
NPMR_NP_theta_4 73.95 74.67 79.22 74.76 100 77.99 77.99 77.46 77.53 78.44 72.19 73.43
NPMR_NP_theta_3 75.18 75.16 75.23 75.15 77.99 100 97.24 78.29 79.1 79.09 73.68 73.67
YK1312_12N_NP_theta 74.59 75.65 76.2 75.69 77.99 97.24 100 79.14 79.12 79.43 73.05 74.54
NPMR_NP_theta_2 73.77 74.62 74.71 74.38 77.46 78.29 79.14 100 93.05 95.89 74.62 76.71
NPMR_NP_theta_1 73.98 74.72 74.36 74.59 77.53 79.1 79.12 93.05 100 96.18 73.08 80.6
NPMR_NP_theta_5 74.38 74.91 75.17 76.24 78.44 79.09 79.43 95.89 96.18 100 72.78 72.48
NPMR_NP_iota_1 72.14 72.86 71.74 71.96 72.19 73.68 73.05 74.62 73.08 72.78 100 99.02
YK1309_1N_NP_iota 72 72.89 72.69 72.33 73.43 73.67 74.54 76.71 80.6 72.48 99.02 100
NP-delta
NP-theta
NP-iota
Percentage
100
95
90
85
80
75

Figure S2

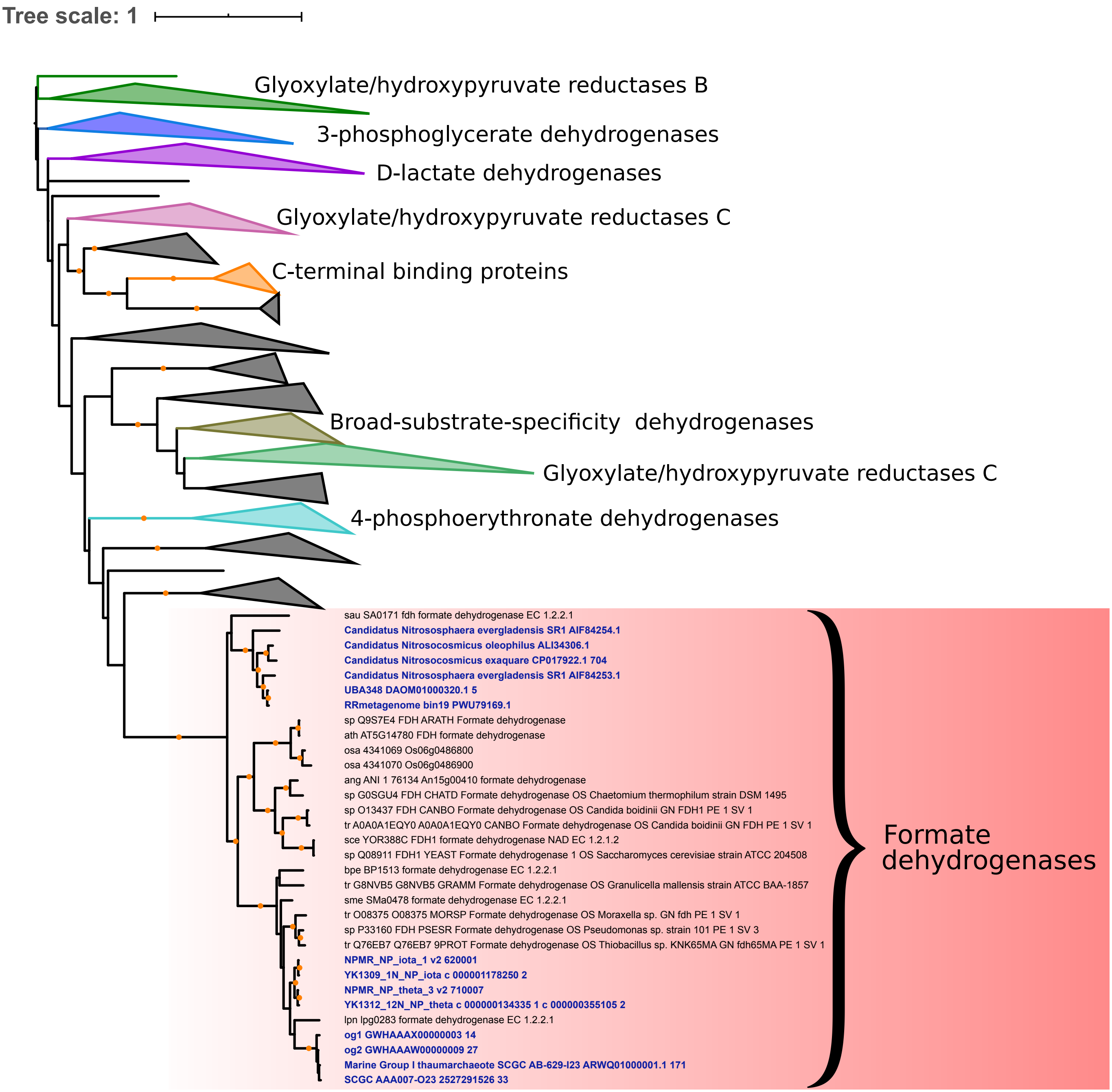

Tree scale: 1
Glyoxylate/hydroxypyruvate reductases B
3-phosphoglycerate dehydrogenases
D-lactate dehydrogenases
Glyoxylate/hydroxypyruvate reductases C
C-terminal binding proteins
Broad-substrate-specificity dehydrogenases
Glyoxylate/hydroxypyruvate reductases C
4-phosphoerythronate dehydrogenases
sau SA0171 fdh formate dehydrogenase EC 1.2.2.1
Candidatus Nitrososphaera evergladensis SR1 AIF84254.1
Candidatus Nitrosocosmicus oleophilus ALI34306.1
Candidatus Nitrosocosmicus exaquare CP017922.1 704
Candidatus Nitrososphaera evergladensis SR1 AIF84253.1
UBA348 DAOM01000320.1 5
RRmetagenome bin19 PWU79169.1
sp Q9S7E4 FDH ARATH Formate dehydrogenase
ath AT5G14780 FDH formate dehydrogenase
osa 4341069 Os06g0486800
osa 4341070 Os06g0486900
ang ANI 1 76134 An15g00410 formate dehydrogenase
sp G0SGU4 FDH CHATD Formate dehydrogenase OS Chaetomium thermophilum strain DSM 1495
sp O13437 FDH CANBO Formate dehydrogenase OS Candida boidinii GN FDH1 PE 1 SV 1
tr A0A0A1EQY0 A0A0A1EQY0 CANBO Formate dehydrogenase OS Candida boidinii GN FDH PE 1 SV 1
sce YOR388C FDH1 formate dehydrogenase NAD EC 1.2.1.2
sp Q08911 FDH1 YEAST Formate dehydrogenase 1 OS Saccharomyces cerevisiae strain ATCC 204508
bpe BP1513 formate dehydrogenase EC 1.2.2.1
tr G8NVB5 G8NVB5 GRAMM Formate dehydrogenase OS Granulicella mallensis strain ATCC BAA-1857
sme SMa0478 formate dehydrogenase EC 1.2.2.1
tr O08375 O08375 MORSP Formate dehydrogenase OS Moraxella sp. GN fdh PE 1 SV 1
sp P33160 FDH PSESR Formate dehydrogenase OS Pseudomonas sp. strain 101 PE 1 SV 3
tr Q76EB7 Q76EB7 9PROT Formate dehydrogenase OS Thiobacillus sp. KNK65MA GN fdh65MA PE 1 SV 1
NPMR_NP_iota_1 v2 620001
YK1309_1N_NP_iota c 000001178250 2
NPMR_NP_theta_3 v2 710007
YK1312_12N_NP_theta c 000000134335 1 c 000000355105 2
lpn lpg0283 formate dehydrogenase EC 1.2.2.1
og1 GWHAAAX00000003 14
og2 GWHAAAW00000009 27
Marine Group I thaumarchaeote SCGC AB-629-I23 ARWQ01000001.1 171
SCGC AAA007-O23 2527291526 33
Formate dehydrogenases


Figure S3

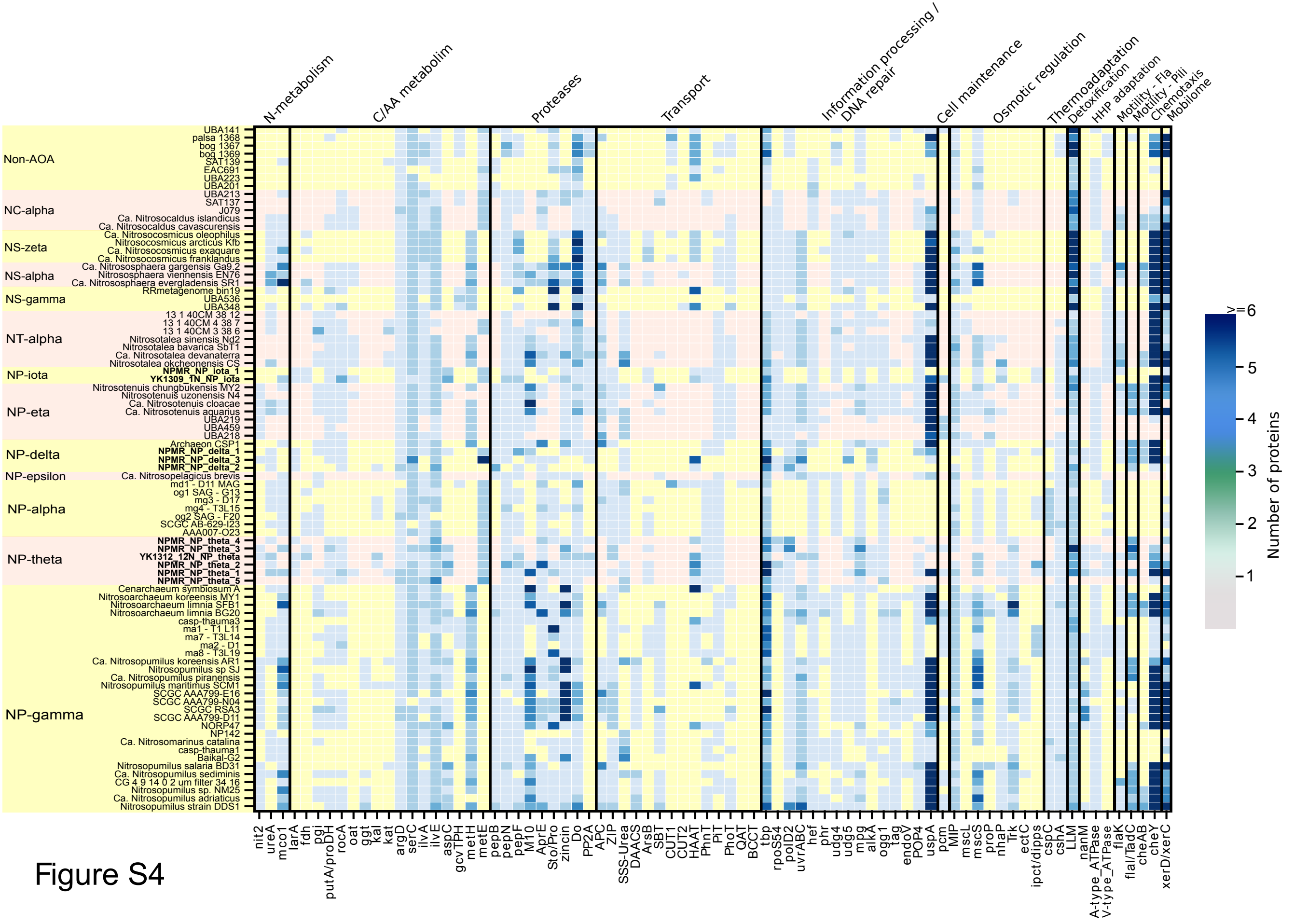
N-metabolism
C/AA metabolim
Proteases
Transport
Information processing / DNA repair
Cell maintenance
Osmotic regulation
Thermoadaptation
Detoxification
HHP adaptation
Motility - Fla
Motility - Pili
Chemotaxis
Mobilome
Non-AOA
NC-alpha
NS-zeta
NS-alpha
NS-gamma
NT-alpha
NP-iota
NP-eta
NP-delta
NP-epsilon
NP-alpha
NP-theta
NP-gamma
Number of proteins
>=6

Figure S4

```
                                                                   54                  68
Gster_LDH           1 MKNNGG--ARVVVIGA-GFVGASYVFALMNQGIADEIVLIDANESKAIGDAM--DFNHGKVFAP--KPVDI-WHGDYDDC
Tth_LDH             1 --------MKVGIVGS-GMVGSATAYALALLGVAREVVLVDLDRKLAQAHAE--DILHATPFA---HPVWVRA-GSYGDL
Blon_LDH            1 MAETTVKPTKLAVIGA-GAVGSTLAFAAAQRGIAREIVLEDIAKERVEAEVL--DMQHGSSFY---PTVSIDGSDDPEIC
Ctep_MDH            1 --------MKITVIGA-GNVGATTAFRLAEKQLARELVLLDVVEGIPQGKAL--DMYESGPVGL--FDTKVTGSNDYADT
Nvie_MDH            1 --------MTITVIGS-GKVGASAALNCGLRELDD-ILLLDIVQGLPQGEAM--DINHQLSERG--SDSVARGSNNYEDM
Nmar_MDH            1 ---------MITIIGS-GKVGGDAALFSALKRLDDQILLLDVAEGLPQGEAM--DINHMLSEQG--IDVEVKGSNNFEDM
Nkor_MDH            1 ---------MITIIGS-GKVGGDAALFSALKRLDDQILLLDVAEGLPQGEAM--DINHMLSEQG--IDVEVKGSSNFEDM
NPMRtheta1_MDH      1 ---------MITIIGA-GKVGGGAALFSALRNLDDQILLLDIVEGLPQGEAM--DINHALSEQG--IDVEIKGSNDYSDM
NPMRtheta5_MDH      1 ---------MITIIGA-GKVGGGAALFSALRNLDDQILLLDIVEGLPQGEAM--DINHALSEQG--IDVEIKGSNDYSDM
NPMRtheta2_MDH      1 ---------VITIIGA-GKVGGDAALFSALRNLDDQILLLDIVKGLPQGEAM--DINHALSEQG--IDVEIKGSNDYSDM
YK1312theta_MDH     1 ---------MITIIGA-GKVGGEAALFSALKKLDDQILLLDIIEGLPQGEAM--DINHMLSEQG--IDVEVKGSNDYSDM
NPMRtheta4_MDH      1 ---------MITIIGA-GKVGGDAALFSALKKLDNQILLLDIIEGLPQGEAM--DINHTLSEQG--IDVDIKGSNDYSDM
Nbrev_MDH           1 ---------MITIIGA-GKVGGDAALFSALRRLDDEILLLDIAEGLPQGEAM--DLNHMLSEQG--IDVNVKGSNNYEDM
NPMRdelta3_MDH      1 ---------MITIIGS-GKVGGAAALFTALKKLDEQILLLDVVKGLPQGEAM--DLNHALSEQG--VDVEIIGSNDYSDM
NPMRdelta2_MDH      1 ---------MITIIGS-GKVGGEAALFTALKKLDDQILLLDIVKGLPQGEAM--DLNHSLSELG--IDVEIIGSNDYSDM
NPMRiota1_MDH       1 ---------MITIVGA-GKVGGAAAISIALRNLSDEILLLDVIKGLPEGEAM--DINHMLSEKG--INVDVRGSNDYSDI
YK1309iota_MDH      1 ---------MITIVGA-GKVGGAAAISIALRNLSDEILLLDVIKGLPEGEAM--DINHMLSEKG--INVDVRGSNDYSDI
Nuzo_MDH            1 ---------MLTIIGS-GKVGGDAALFSALRKVDKDILLLDVVNGLPQGEAM--DLNHMLSEQG--VDVHIRGSNDYADM
Mjan_MDH            1 --------MKVTIIGASGRVGSATALLLAKEPFMKDLVLIGREHSINKLEGLREDIYDALAGTRSDANIYVESDENLRII
Iisl_MDH            1 MARI---PYKVAVIGT-GRVGATFAYTMAVVPGIARMTLVDVVPGLAKGVME--DIKHAAAVFR--RSITVEAFEDVSKV
Msed_MDH            1 M-------AKVGFIGA-GKIGQTIAYSALVSGAVDEAVIYDIIPELPDKFEH--ELRHAFATKG--IKANVLGTNSLDDV

                                     102
Gster_LDH          73 RDADLVVICAGANQKPGETRLDLVDKNIAIFRSIVESVMASGFQGLFLVATNPVDILTYATWKFSGLPHERVIGSGTILD
Tth_LDH            66 EGARAVVLAAGVAQRPGETRLQLLDRNAQVFAQVVPRVLEAAPEAVLLVATNPVDVMTQVAYRLSGLPPGRVVGSGTILD
Blon_LDH           75 RDADMVVITAGPRQKPGQSRLELVGATVNILKAIMPNLVKVAPNAIYMLITNPVDIATHVAQKLTGLPENQIFGSGTNLD
Ctep_MDH           68 ANSDIVVITAGLPRKPGMTREDLLSMNAGIVREVTGRIMEHSKNPIIVVVSNPLDIMTHVAWQKSGLPKERVIGMAGVLD
Nvie_MDH           67 RGSDYVVLVAGVGRKPGMTRMDLLKINAGIVKDVASKIATYAKDATVIVVTNPLDPMTYLALKTIGAQKSKVMGMGGMLD
Nmar_MDH           67 KGSNIVVVVAGSGRKPGMTRMDLLKINASIVKSVVENVKKYADDSMIIPVTNPLDPMAYITYKVSGFDRSRVFGMGGMLD
Nkor_MDH           67 KGSKIIVVVAGSGRKPGMTRMDLLKINATIVKSVVENIKKYANDSMIIPVTNPLDPMAYITYKVSGFDRSRVFGMGGMLD
NPMRtheta1_MDH     67 KGSNIVVVVAGLGRKPGMTRMDLLQKNAGIVKSVVENIKKYADNSMIIPVTNPLDPMTYICYKVSGFERNRVFGMGGMLD
NPMRtheta5_MDH     67 KGSNIVVVVAGLGRKPGMTRMDLLQKNAGIVKSVVENIKKYADDSMIIPVTNPLDPMAYICYKVSGFERNRVFGMGGMLD
NPMRtheta2_MDH     67 KGSNIVVVVAGLGRKPGMTRMDLLQKNAGIVKSVVENIKKYADDSIIIPVTNPLDPMTYICYKVSGFERNRVFGMGGMLD
YK1312theta_MDH    67 KGSDIVVTVAGSGRKPGMTRMDLLKINAGVVKSVVENIKKYANDAMIIPVTNPLDPMAYICYKVSGFEKNRIFGMGGMLD
NPMRtheta4_MDH     67 KGSDIVVVVAGLGRKPGMTRMDLLQKNAGIVKSVVENIKKYANDSMIIPVTNPLDPMAHICYKVSGFEKNRVIGMGGMLD
Nbrev_MDH          67 KGSKIVVVVAGAGRKPGMTRMDLLKINAGVVKDVVGNIKKFADDAMIIPVTNPLDPMAQIAYKTSGFDRSRVFGMGGMLD
NPMRdelta3_MDH     67 KDSKIVVIVAGSGRKPGMTRMDLLKINAGVVKDVVENIKKYSNDAILVPVTNPLDPMAHIAYKTSGFERNRILGMGNMLD
NPMRdelta2_MDH     67 KNSKIVVVVAGSGRKPGMTRMDLLKINAGVVKDVVENIKKYSKDSFIIPVTNPLDPMAHIAYKVSGFSRNRIFGMGNMLD
NPMRiota1_MDH      67 KGSEIVVVVAGSGRKPGMTRMDLLKVNASVVKDVTENIKKYANNSIIVPVTNPLDPMAYVAFKVSGFSKNRIVGMGNMLD
YK1309iota_MDH     67 KGSEIVVVVAGSGRKPGMTRMDLLKVNASVVKDVTENIKKYANNSIIVPVTNPLDPMAYVAFKVSGFSKNRIVGMGNMLD
Nuzo_MDH           67 KGSDIVVVVAGSGRKPGMTRMDLLKINAGIVKDVVENIKKHASNAIIVPVTNPLDPMVHITYKTSGFEPSRIVGMGNMLD
Mjan_MDH           73 DESDVVIITSGVPRKEGMSRMDLAKTNAKIVGKYAKKIAEICDTKIF-VITNPVDVMTYKALVDSKFERNQVFGLGTHLD
Iisl_MDH           73 ENADAIVITAGKPRKADMSRRDLANVNAQIIRDIGDKLRDRNPGALYVVVTNPVDVMTMVLDDVIG-SKGTVIGTGTSLD
Msed_MDH           69 SGMDIVVISAGKPRKPGMSRRDLFVDNAKIMIDLAQKLPSKNPGAIYLMVANPVDMMASVFMKY---SKQFTISAGDQVE

                        171                        199
Gster_LDH         153 TARFRFLLGEYFSVAPQNVHAYIIGEHGDTELPVWSQAYIGVMPIRKLVESKGEE--AQKDLERIFVNVRDAAYQIIE--
Tth_LDH           146 TARFRALLAEYLRVAPQSVHAYVLGEHGDSEVLVWSSAQVGGVPLLEFAEARGRA-LSPEDRARIDEGVRRAAYRIIE--
Blon_LDH          155 SARLRFLIAQQTGVNVKNVHAYIAGEHGDSEVPLWESATIGGVPMCDWTPLPGHDPLDADKREEIHQEVKNAAYKIIN--
Ctep_MDH          148 SARFRSFIAMELGVSMQDVTACVLGGHGDAMVPVVKYTTVAGIPVADL--------ISAERIAELVERTRTGGAEIVNHL
Nvie_MDH          147 LSRFKSYIQEATGVSRDSIQAMVISEHGENMLPLTRFSSLGGIPLHDF--------ITKEQATDIFEKTKKVAAEVIA--
Nmar_MDH          147 LSRFRQFIHEATGHSRDSIRALVIGEHGENMLPLPRFSSVSGIPLPSL--------LPKEKLEELVQNTKQVAAKVIE--
Nkor_MDH          147 LSRFRQFIHEATGHSRDSIRALVIGEHGENMLPLPRFSSVSGIPLSSF--------LPKQKLDELVQNTKQVAAKVIE--
NPMRtheta1_MDH    147 ISRFKQFIHEATGHSRHSIRALVIGEHGENMLPLPRFSSVSGIPLTSL--------LSKEKLDELVQNTRNVAAKVIE--
NPMRtheta5_MDH    147 ISRFKQFIHEATGHSRHSIRALVIGEHGENMLPLPRFSSVSGIPLTSL--------LPKEKLDELVQNTRQVAAKVIE--
NPMRtheta2_MDH    147 LSRFKQFIHEATGHSRHSIRALVIGEHGENMLPLPRFSSVSGIPLTSL--------LSKEKIDELVQNTRQVAAKVIE--
YK1312theta_MDH   147 ISRFIQFIHEATGHSRDSIRALVIGEHGENMLPLPRFSSVSGIPLTSL--------LSKEKLDELVQNTKQVAAKVIE--
NPMRtheta4_MDH    147 LSRFKQFIHEATGYSRDSIRALVIGEHGENMLPLPRFSSVAGIPLVSL--------LSKEKLDELIQNTRQIAAKVIE--
Nbrev_MDH         147 LSRFKQFIHEATGYSRESTKALVIGEHGENMLPLTRFATVSGIPLPTL--------LPKEKLDEIFTATKGVAAEVIK--
NPMRdelta3_MDH    147 LSRFSQFIHEATGQSRESIRALVIGEHGENMLPLIRFSSVSGIPLTSL--------LAKDKLEELEKNTRQVAAKVIE--
NPMRdelta2_MDH    147 ISRFTQFIHEATGYSRQSIRALVIGEHGENMLPLMRFSSVSGIPLTSL--------LSKEKLDELEKNTRQVAAKVIE--
NPMRiota1_MDH     147 LSRFKQFIHESTGFSRDSIRALVIGEHGENMVPLPRFSSVAGIPLMSL--------LSKEKIDELVVSTREVAAKVIE--
YK1309iota_MDH    147 LSRFKQFIHESTGFSRDSIRALVIGEHGENMVPLPRFSSVAGIPLMSL--------LPKEKIDELVISTREVAAKVIE--
Nuzo_MDH          147 LSRFIQFIHESTGHSRDSIRALVIGEHGENMLPLPRFSTVSGIPLASI--------LPKEKLDQIVKDTRGVAAKVIE--
Mjan_MDH          152 SLRFKVAIAKFFGVHIDEVRTRIIGEHGDSMVPLLSATSIGGIPIQKFER------FKELPIDEIIEDVKTKGEQIIR--
Iisl_MDH          152 TFRFRAAVSELLNVPIVAVDGYVVGEHGEEAFVAWSTVTIKGIHIDQYIKERNIN-ISR---EQIEKYVKDVAASIIA--
Msed_MDH          146 TMRMRSFIAKKLKIPVTSVDGFVGGEHGEDAVVLWSTVKIKGKPVDEFN-------INK---DEVSDYVKKIPGEIIR--
```

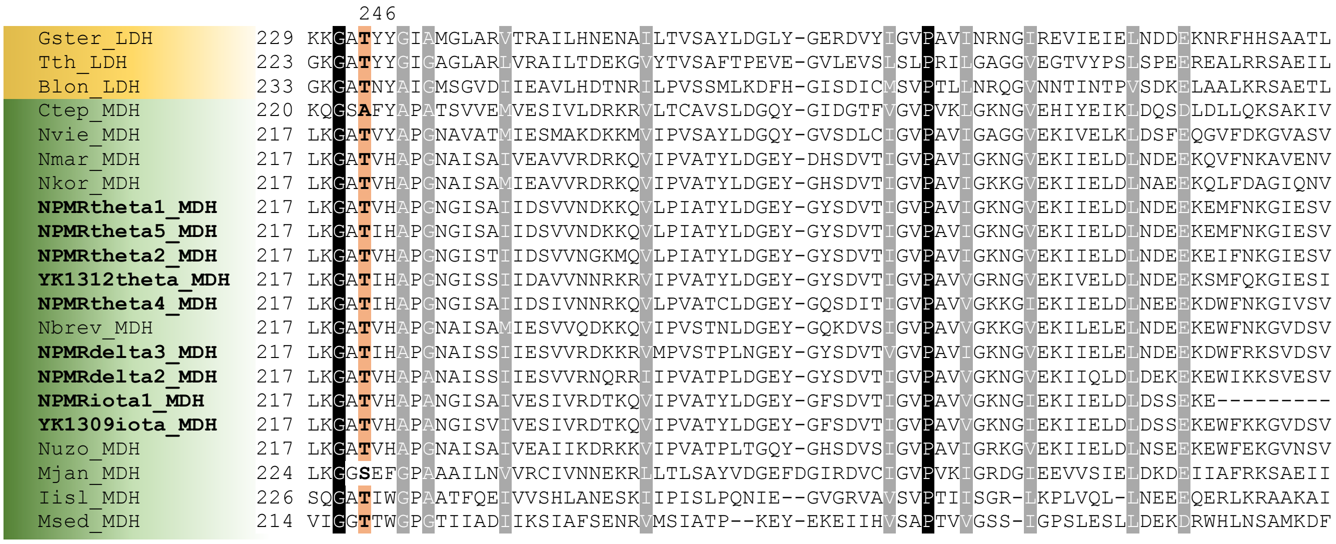


**Fig. S5.** Sequence alignment of lactate dehydrogenase (LDH) and malate dehydrogenase (MDH) homologs, shaded in yellow and green, respectively. Conserved residues are shaded in black and grey. The universally conserved in the LDH/MDH superfamily substrate binding Arg171 is indicated in red. Residues important for substrate discrimination and active site architecture are shaded in orange and discussed in the text. The residue determining the cofactor specificity is shaded in green. Residue numbering refers to the LDH numbering as in Roche *et al.* 2019.

Primary sequences from the following organisms were used to generate the alignment (Uniprot accession numbers in parentheses): Gster, *Geobacillus stearothermophilus* (P00344); Tth, *Thermus thermophilus* (Q5SJA1); Blon, *Bifidobacterium longum* (E8ME30); Ctep, *Chlorobaculum tepidum* (P80039); Nvie, *Nitrososphaera viennensis* (A0A060HG74); Nmar, *Nitrosopumilus maritimus* (A9A450); Nkor, *Nitrosarchaeum koreense* (F9CUM5); Nbrev, *Ca.* Nitrosopelagicus brevis (A0A0A7V4F4); Nuzo, *Ca.* Nitrosotenuis uzonensis (V6AR53); Mjan, *Methanocaldococcus jannaschii* (Q60176); Iisl, *Ignicoccus islandicus* (A0A0U3FQH7); Msed, *Metallosphaera sedula* (A4YDY0). Sequences from the marine sediment AOA MAGs reported in this study are in bold and their locus tags can be found in Table S3.

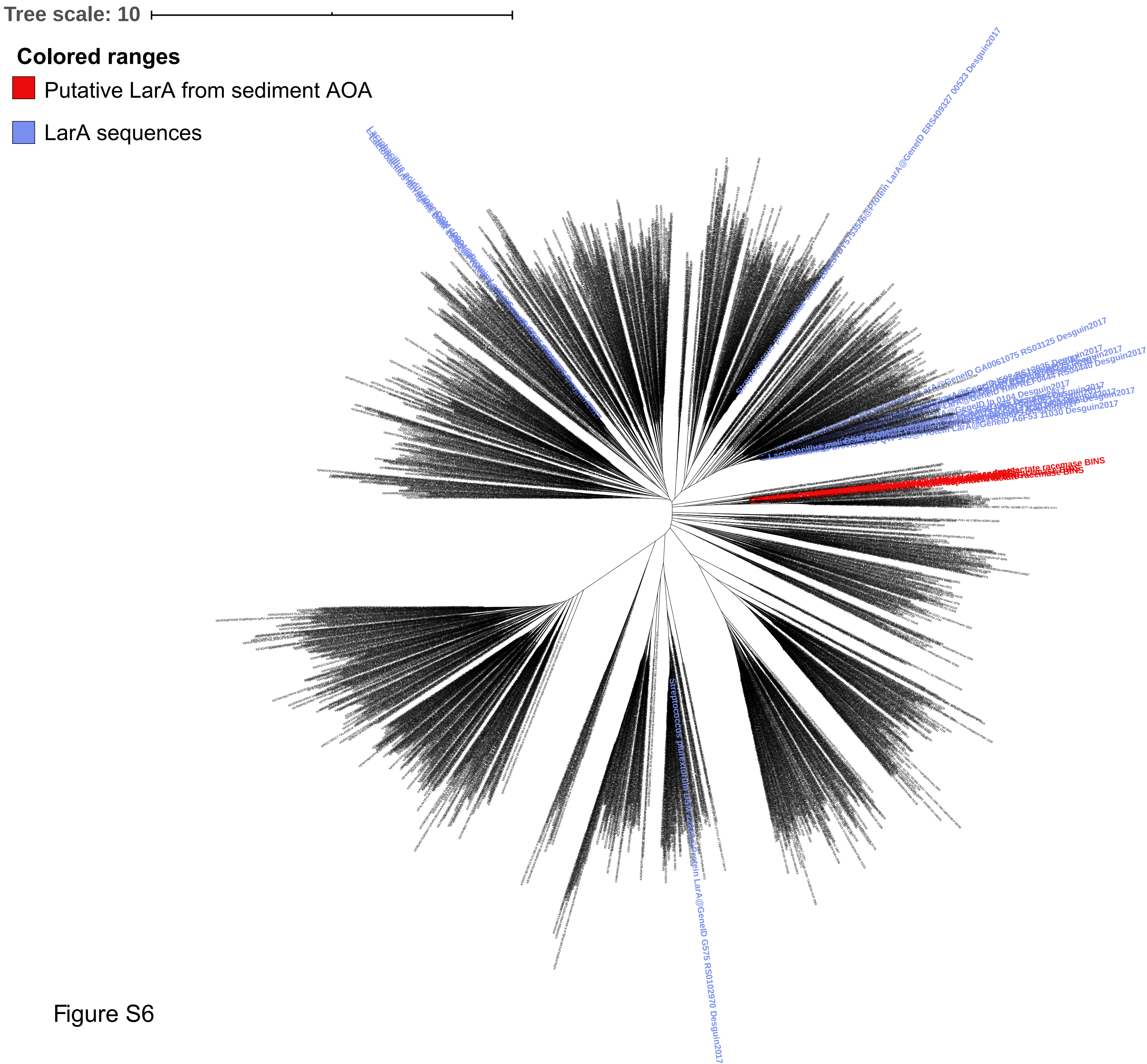

Tree scale: 10
Colored ranges
Putative LarA from sediment AOA
LarA sequences


Figure S6
